# Towards self-regeneration: exploring the limits of protein synthesis in the PURE cell-free transcription-translation system

**DOI:** 10.1101/2024.04.03.587879

**Authors:** Ragunathan Bava Ganesh, Sebastian J. Maerkl

## Abstract

Self-regeneration is a key function of living systems that needs to be recapitulated *in vitro* to create a living synthetic cell. A major limiting factor for protein self-regeneration in the PURE cell-free transcription-translation system is its high protein concentration, which far exceed the system’s protein synthesis rate. Here we were able to drastically reduce the non-ribosomal PURE protein concentration up to 97.3% while increasing protein synthesis efficiency. Although crowding agents were not effective in the original PURE formulation, we found that in highly dilute PURE formulations addition of 6% dextran considerably increased protein synthesis rate and total protein yield. These new PURE formulations will be useful for many cell-free synthetic biology applications and we estimate that PURE can now support the complete self-regeneration of all 36 non-ribosomal proteins, which is a critical step towards the development of a universal biochemical constructor and living synthetic cell.

## Introduction

One of the goals of bottom-up synthetic biology is to build synthetic cells. Self-regeneration, evolvability, adaptability, information processing, and metabolism are some of the main functions of living systems^1,2,3,4^. Various research groups re-created some of these functions in cell-free environments by implementing DNA replication^5,6^, metabolism^7,8^, protein regeneration^9^, and compartmentalization^10,11^. Cell-free systems are ideal starting points for the bottom-up re-creation of fundamental cellular processes. Particularly the PURE (Protein synthesis Using Recombinant Elements) recombinant cell-free system is an excellent platform for engineering purposes, because the composition and components of the system are defined and can be manipulated at will^12^. The PURE system consists of 36 non-ribosomal proteins for transcription (1 protein), translation (10 proteins), aminoacylation (20 proteins), and energy regeneration (5 proteins) (Fig 1a). To create the PURE system, all 36 non-ribosomal proteins are expressed and purified after which they are combined with ribosomes, tRNAs, and an energy mix containing the necessary small molecule components. To simplify this process we recently developed a facile process that co-expresses and co-purifies all 36 non-ribosomal proteins, drastically reducing the labor and cost associated with PURE production^13^.

**Figure 1:**
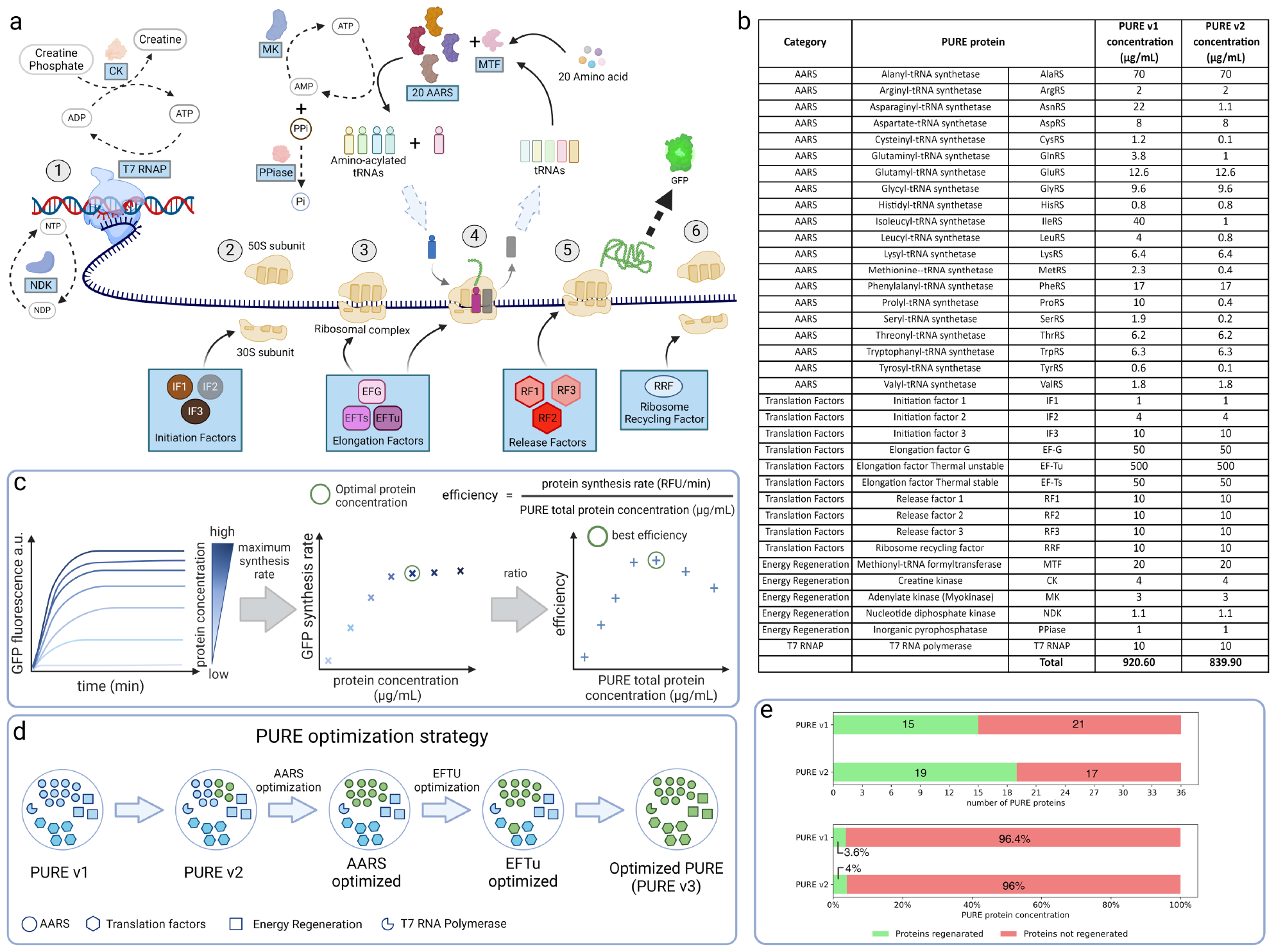
The PURE system comprises ribosomes and 36 proteins for transcription, translation, aminoacylation, and energy regeneration. a) A schematic of all 36 PURE proteins and their functions. b) A list of all 36 proteins and their concentrations in the original PURE v1 and PURE v2 system formulations. c) Schematic representation of the optimization procedure and criteria for selecting the optimal protein concentrations. d) Schematic representation of the optimization strategy showing the order in which the PURE proteins were optimized. e) Estimates of the current self-regenerative capacity of the PURE system, showing the number of proteins and the total PURE non-ribosomal protein concentration that can theoretically be self-regenerated in a chemostat environment.

To build a synthetic cell, key cellular components such as DNA^5,6^, proteins^9,14,15^, ribosomes^16,17,18,19^, tRNAs^20^, and lipids^21^ will likely need to be regenerated. So far, attempts are mainly focusing on achieving regeneration of these individually, although in some cases attempts were made to couple protein regeneration and DNA replication^22,23^. The regeneration potential of all 36 PURE proteins was assessed individually and summarized in a “pureiodic table”^24^. We previously were able to demonstrate protein self-regeneration in the PURE system using a microfluidic chemostat, achieving continuous and sustained self-regeneration of up to 7 PURE proteins^9^. This work showed that it is possible to self-regenerate essential PURE proteins *in vitro*, but also indicated that self-regenerating all 36 non-ribosomal proteins is a major challenge due to the limited synthesis capacity of the PURE system, which we estimated to be at least one order of magnitude too low for complete self-regeneration of all non-ribosomal PURE proteins. We estimated that the original PURE system could self-regenerate up to 15 (42%) of the 36 proteins, which corresponds to only 3.6% of the total non-ribosomal protein content. We recently explored the possibility to decrease chemostat dilution rates. By decreasing dilution rates less protein needed to be synthesized per unit time. Unfortunately, simply decreasing the dilution rate was counterproductive, as synthesis rates decreased significantly. To overcome this problem, we developed a next-generation chemostat that incorporated a dialysis membrane for continuous diffusion of small molecules in and out of the chemostat^25^. With this system dilution rates could be drastically decreased without negatively impacting synthesis rates.

Decreased dilution rates are by themselves likely not sufficient to achieve complete non-ribosomal protein self-regeneration in the PURE system. We therefore turned our attention to optimizing the PURE system itself. Previous approaches to optimizing the PURE system focused on increasing total synthesis yield^26,27,28^. This was achieved by either increasing the concentration of specific PURE proteins or introducing new proteins to the system. Although these approaches achieved higher protein yields, they are not viable solutions for solving the problem of self-regenerative capacity, as increasing protein concentration or adding proteins increases the total protein concentration that needs to be self-regenerated. It is therefore the ratio of protein synthesis rate to total protein content of the PURE system, which we call protein synthesis efficiency, that we set out to improve.

We previously reported a modest reduction of total non-ribosomal protein concentration from 920.6 µg/mL (PURE v1) to 839.90 µg/mL (PURE v2) by optimizing the concentration of 9 proteins without negatively impacting protein synthesis rate^9^. This led to a projected improvement in self-regeneration capacity of 19 (53%) instead of 15 (42%) proteins. But we wondered whether it might be possible to drastically reduce the total non-ribosomal protein concentration of the PURE system. Although this might lead to a concurrent decrease in protein synthesis rate, it might ultimately lead to an overall improvement as long as the ratio of protein synthesis rate to total protein concentration increases. We therefore set out to explore the limits of the PURE system by systematically characterizing and minimizing the concentration of every non-ribosomal protein and arrived at drastically reduced PURE formulations with increased protein synthesis efficiency.

## Results

### PURE optimization

The primary objective was to systematically reduce overall PURE non-ribosomal protein concentration while maintaining or improving protein synthesis efficiency. We define protein synthesis efficiency as the ratio of protein synthesis rate to total non-ribosomal protein concentration. The 36 PURE proteins were divided into 4 subsets for optimization: aminoacyl-tRNA synthetases (AARS) (20 proteins), initiation and release factors (IRFs) (7 proteins), elongation factors (EFs) (3 proteins), and T7 RNAP + energy regeneration (ER) (6 proteins) (Fig 1a,b).

The optimization strategy was to titrate and optimize each protein individually (Fig 1c). From previous work, we hypothesized that most, if not all, protein titrations would result in a synthesis rate that saturates above a critical protein concentration. We thus would choose a protein concentration near this critical threshold at which protein synthesis efficiency is optimal. Our optimization started with the previously optimized PURE v2 formulation (Fig 1b) by optimizing all 20 AARSs after which we optimized EF-Tu which is by far the most abundant non-ribosomal protein in the PURE system. After the AARSs and EF-Tu were optimized, we used this PURE formulation as the base for optimizing the remaining three categories: initiation and release factors, elongation factors (minus EF-Tu), and the T7 RNAP + energy regeneration enzymes (Fig 1d).

When titrating individual proteins, we predominantly observed monotonically saturating curves (Figure 2 a-d, and Supp Fig 1 - 6). Most proteins were found to be essential, although several proteins were found to be non-essential. Some of these non-essential proteins were expected to be non-essential whereas others should have been essential. The latter are probably best explained by minor contaminations of these proteins which are sufficient for function (Supp Fig 7). The AARS subset was reduced by 93.2% from 226.5 µg/mL to 15.31 µg/mL (Fig 2e). The IRF subset was reduced 79.5% from 55 µg/mL to 11.25 µg/mL (Fig 2f), the T7 RNAP + ER subset was reduced 74.4% from 39.1 µg/mL to 10 µg/mL (Fig 2g), and the EFs subset concentration was reduced 72.5% from 600 µg/mL to 165 µg/mL (Fig 2h).

**Figure 2:**
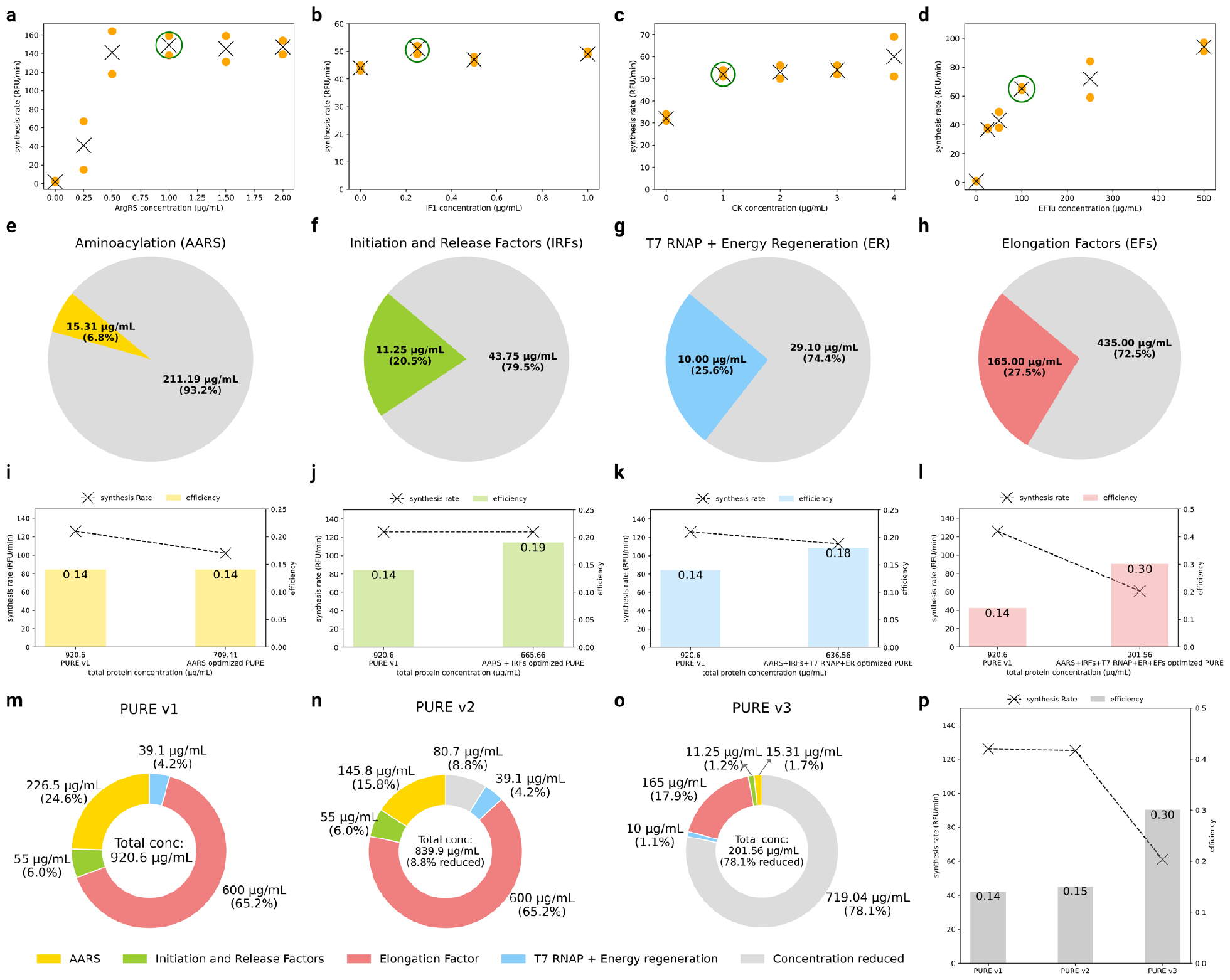
PURE optimization summary. (a-d) Examples of the titration experiments. Shown are the results for ArgRS (a), IF1 (b), CK (c), and EFTu (d) as examples for each of the 4 categories: aminoacylation, initiation and release factors, T7 RNAP + energy regeneration and elongation factors. Titrations of all 36 non-ribosomal proteins are provided in the supplement (Supp Fig. 1 - 6). (e-h) The original and optimized concentration of each subset and the percent reduction compared to the original PURE v1 system. (i-l) The result of the swap-in experiment in which the optimized sub-set concentrations were used instead and compared to the original PURE v1 formulation. Line graphs show the average synthesis rate and the bar plots indicate the average efficiency of the PURE system (n=2). (m-o) The composition of different PURE versions with the percent reduction compared to the PURE v1 system. (p) The average synthesis rate and average efficiency of all 3 PURE versions (n=2). Line graphs show the average synthesis rate and the bar plots indicate the average efficiency of the PURE system.

We then consecutively combined the various reduced subsets to derive a new PURE system. Despite the relatively large reduction in protein concentration in the AARS subset, due to the concurrent reduction in protein synthesis rate, reducing AARSs led to protein synthesis efficiency remaining unchanged at 0.14 (Fig 2i). We then added the IRF-optimized subset which did not impact protein synthesis rate and thus led to a slight increase in protein synthesis efficiency to 0.19 (Fig 2j). Addition of the T7 RNAP + ER optimized subset was similar to the AARS subset in that it slightly reduced the protein synthesis rate while maintaining a protein synthesis efficiency of 0.18 (Fig 2k). Only when adding the EF subset did we observe a major decrease in protein synthesis rate to roughly half the rate of the original PURE recipe. Despite this decrease in synthesis rate, protein synthesis efficiency increased to 0.30 due to the large reduction in protein concentration in this subset (Fig 2l). We termed the resulting PURE formulation PURE v3.

Compared to the original PURE formulation (PURE v1) we were able to reduce non-ribosomal protein concentration from 920.6 µg/mL to 201.56 µg/mL, which is a reduction of 719.04 µg/mL or 78.1% (Fig 2 m-o). Protein synthesis rate decreased from 126 RFU/min to 61 RFU/min while protein synthesis efficiency was increased two-fold from 0.14 to 0.30 (Fig 2p).

### Exploring the limits of PURE transcription and translation

Motivated by this result, we were curious to explore the limits of PURE cell-free protein synthesis *in vitro*. We diluted PURE v3 to the following concentrations: 90 µg/mL (PURE v3.90), 50 µg/mL (PURE v3.50), 25 µg/mL (PURE v3.25), and 10 µg/mL (PURE v3.10) (Fig 3a, and Supp Table 1,2). To put these concentrations in perspective, the total protein concentration of *E. coli* is estimated to be 200-320 mg/mL^29^. In comparison to *E. coli*, PURE v1 non-ribosomal protein concentration is 217-348 times lower; PURE v3 is diluted by four orders of magnitude and PURE v3.10 diluted by five orders of magnitude. For testing protein expression using these diluted PURE formulations we used commercial NEB *E. coli* ribosomes as opposed to his-tag purified ribosomes which were used in the previous experiments because commercial NEB ribosomes generally give rise to higher synthesis rates as compared to home-made his-tagged ribosomes.

**Figure 3:**
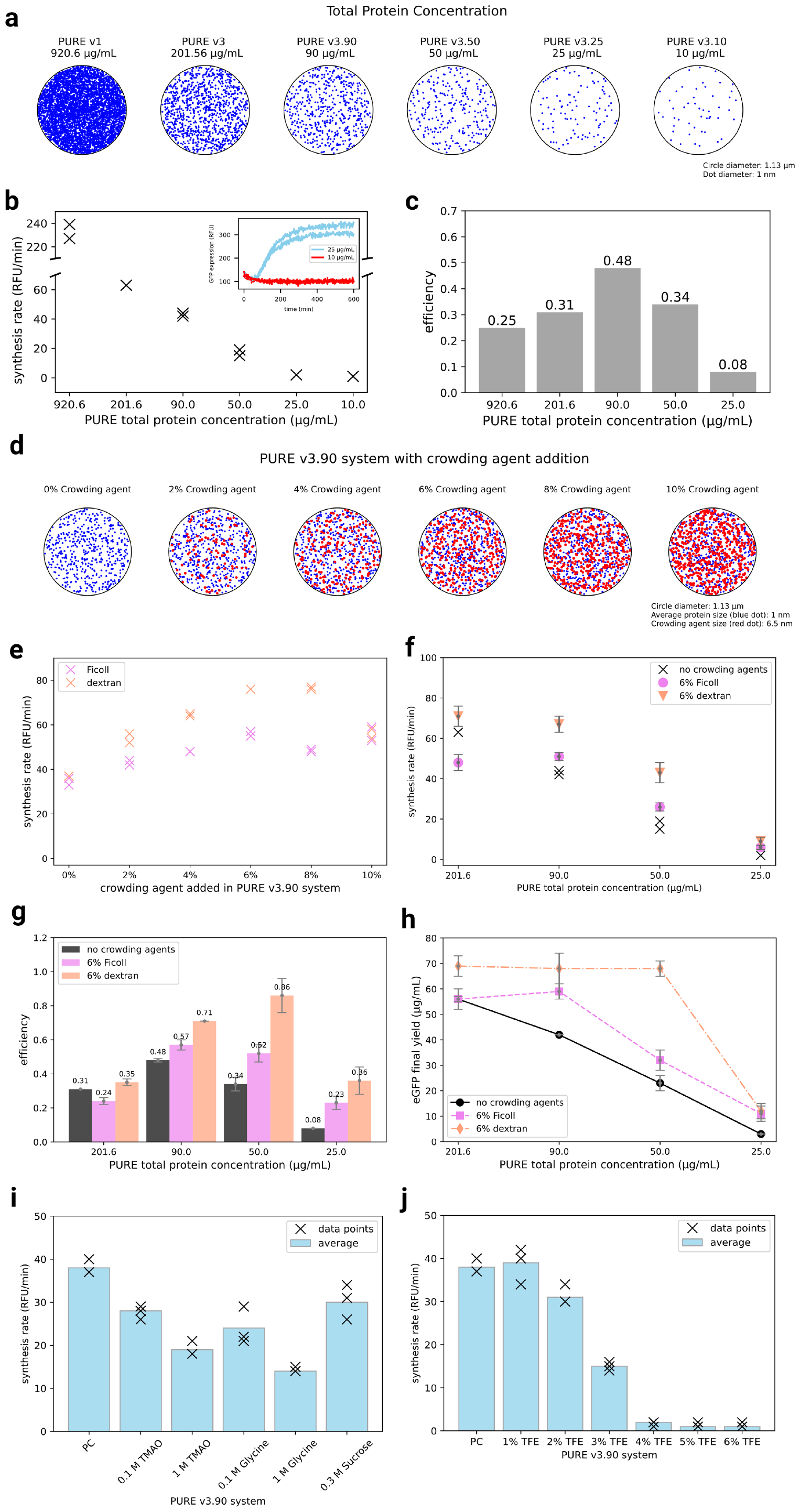
Exploring the concentration limits of the PURE system and the role of crowding agents. (a) A schematic of the number of PURE proteins present in a 2 µm^3^ volume with a diameter of 1.13 µm and height of 2 µm for the original PURE v1 to the ultra-diluted PURE v3.10. Each blue dot represents a protein. (b) Synthesis rate of various PURE systems plotted against the total protein concentration present in that particular PURE system formulation. The inset shows the real-time synthesis curves for the 25 µg/mL and the 10 µg/mL PURE system (n=2). (c) Average protein synthesis efficiency of the various PURE systems. PURE v3.90 achieved the highest protein synthesis efficiency of 0.48, which is almost double that of the original PUREv1 formulation (n=2). (d) A schematic showing the concentration of crowding agent (red dots) in relation to the non-ribosomal protein concentration (blue dots) in the PURE v3.90 system. (e) Crowding agent titration in PURE v3.90 system. The synthesis rate is plotted against the final concentration of crowding agent (n=2). (f) The synthesis rate of diluted PURE systems with and without crowding agent. (g) The efficiency of diluted PURE systems with and without crowding agent. (h) The final protein yield of diluted PURE systems with and without crowding agent (n=2 for PURE without crowding agents, n=4 for PURE with crowding agents except PURE v3.90 + 6% dextran, and n=10 for PURE v3.90 + 6% dextran group). Error bar presents mean ± standard deviation. (i) Testing of other potential small molecule crowding agents: trimethylamine N-oxide (TMAO) (0.1 M and 1 M), glycine (0.1 M and 1 M) and sucrose (0.3 M) in PURE v3.90 system (n=3). (j) Effect of crowding agent 2,2,2-trifluoroethanol 1% - 6% (v/v) in PURE v3.90 system (n=3). The black cross (x) shows individual data points and bar plots indicate the average synthesis rate of the PURE system.

PURE v1, PURE v3, PURE v3.90, PURE v3.50, and PURE v3.25 had detectable protein expression, whereas PURE v3.10 didn’t give rise to any protein synthesis (Fig 3b, and Supp Fig 8). The breakpoint for protein synthesis of the PURE system therefore lies somewhere between 10 µg/mL and 25 µg/mL of total non-ribosomal protein concentration. These concentrations are 37 to 92-fold lower than the original PURE system and 8,000 to 20,000-fold lower than cellular protein concentrations. PURE v1 with NEB ribosomes achieved a synthesis rate of 233 RFU/min. PURE v3, with a 4.5-fold reduction in total protein concentration, had a synthesis rate of 63 RFU/min (3.7-fold reduction from v1). PURE v3.90, with an order of magnitude lower protein concentration compared to PURE v1, had a synthesis rate of 43 RFU/min (5.4-fold reduction from v1) and PURE v3.50, with an 18.4-fold reduction in total protein concentration, had a synthesis rate of 17 RFU/min (13.7-fold reduction from v1).

With the use of NEB ribosomes in PURE v1 synthesis rate increased from 126 RFU/min (Fig 2p) to 233 RFU/min. However, PURE v3 synthesis rate did not increase and remained at 63 RFU/min despite using NEB ribosomes, indicating that in these lower concentration formulations something else is likely becoming limiting. To determine whether T7 RNAP could be the limiting factor, we tested new PURE formulations with T7 RNAP at the higher nominal concentration of 10 µg/mL (v3 T7 RNAP full, v3.90 T7 RNAP full, v3.50 T7 RNAP full, v3.25 T7 RNAP full, v3.10 T7 RNAP full) but the nominal T7 RNAP concentration did not increase protein synthesis and even had a negative effect on the system (Supp Fig 9).

In terms of efficiency, both PURE v3.90 and PURE v3.50 had higher efficiency values of 0.48 and 0.34 respectively when compared to the efficiency of 0.31 for the PURE v3 system, and 0.25 for PURE v1 with NEB ribosomes (Fig 3c). Efficiency peaked at a concentration of 90 µg/mL (PURE v3.90) and any further dilutions led to a decrease in efficiency. These results indicate that PURE protein concentration could be further optimized in terms of efficiency and also be diluted drastically by at least 40-fold and still result in protein synthesis.

### Effects of crowding agents

Macromolecular crowding is known to affect various biological processes in the cellular environment^30,31^. Therefore the effect of crowding agents has also been explored in cell-free systems with the hope that inducing molecular crowding could enhance protein synthesis rates or yields^32,33,34^. Commonly used crowding agents include dextran, Ficoll, and polyethylene glycol (PEG). In the PURE system even proteins such as bovine serum albumin (BSA) were used as a crowding agent resulting in a ∼70% increase in protein expression^27^. However, we were interested in non-protein crowding agents to avoid increasing the total protein concentration present in the PURE system. Previous additions of non-protein crowding agents to the PURE system were reported to have both positive and negative effects on performance. PEG-6000 (molecular weight: 6kDa) addition at 0.05 – 0.2% (w/v) had a neutral to negative effect on protein yield^27^. Similarly, 5 – 30% (w/v) Ficoll-70 addition to PURExpress strongly repressed protein expression^35^. On the other hand, addition of 1 – 2.5% (w/v) Ficoll-70 or Ficoll-400 improved protein expression by 26 – 33%^36^. Despite the fact that in our hands crowding agents had mixed results when added to the standard PURE system, we nonetheless decided to re-evaluate crowding agents in the ultra-diluted PURE formulations.

Since the PURE v3.90 system had the highest protein synthesis efficiency, we chose this version for testing crowding agents. The crowding agents Ficoll-70 (∼70 kDa molecular weight) and dextran-70 (∼70 kDa molecular weight) were titrated so that their final concentration in the PURE system ranged from 0% to 10% (Fig 3d). The PURE reactions showed an increase in synthesis rate as a function of crowding agent concentration (Fig 3e). Dextran addition resulted in a larger enhancement in synthesis rate compared to Ficoll at almost all tested concentrations. Only the highest concentration of 10% had equivalent enhancement for both dextran and Ficoll. Addition of dextran almost doubled the synthesis rate from 37 RFU/min to 76 RFU/min and 77 RFU/min at 6% and 8%, respectively. In the case of Ficoll, the increase in synthesis rate was not as high as dextran with the maximum being 54 RFU/min for 6% and 10% Ficoll.

Next, we explored what effect 6% Ficoll or dextran had in PURE v1, v3, v3.90, v3.50, and v3.25 (Fig 3f, Supp Fig 10 - 12). For the standard PURE v1 system, both Ficoll and dextran addition decreased the synthesis rate from 233 RFU/min to 203 RFU/min (-13%) and 197 RFU/min (-15%), respectively. In PURE v3 with a synthesis rate of 63 RFU/min, crowding agents slightly increased the synthesis rate to 71 RFU/min (+13%) and Ficoll decreased synthesis rate to 48 RFU/min (-24%). In the concentrated systems, crowding agents therefore had no major effect. On the contrary, in diluted PURE versions both Ficoll and dextran enhanced synthesis rates. In PURE v3.90, PURE v3.50, and PURE v3.25, dextran addition had a larger enhancement than Ficoll addition. In the PURE v3.90 system, the average synthesis rate increased from 43 RFU/min to 64 RFU/min (+49%) with dextran and to 51 RFU/min (+19%) with Ficoll. For PURE v3.50, dextran addition more than doubled the synthesis rate from 17 RFU/min to 43 RFU/min (+153%), while Ficoll increased the synthesis rate to 26 RFU/min (+53%). PURE v3.25 has a low baseline synthesis rate of 2 RFU/min. Dextran addition increased the synthesis rate to 9 RFU/min while Ficoll addition increased to 6 RFU/min. In summary, previous reports on crowding agent addition showed contradicting results with positive, neutral, and negative effects on a standard PURE system. In our study, Ficoll addition had a negative effect on concentrated PURE systems (PURE v1 and v3) whereas dextran had a negative effect on PURE v1 but had a positive effect on PURE v3. This highlights again the relative ineffectiveness of crowding agents in concentrated PURE systems, but in diluted PURE systems crowding agents have a consistent and considerable effect, improving synthesis rates up to 153%.

PURE v3.50 system had the highest improvement with 6% dextran addition. To determine the optimal crowding agent concentration, we titrated dextran and Ficoll in PURE v3.50 system (Supp Fig 13). A slight increase in synthesis rate was observed when Ficoll concentration increased from 0% to 6% followed by a slight decrease for 8% and 10% Ficoll addition. The largest improvement was observed for 6% Ficoll with synthesis rate increasing from 21 RFU/min to 28 RFU/min. As with PURE v3.90, dextran addition improved synthesis rate more than Ficoll in the PURE v3.50 system, giving rise to an increase in synthesis rate from 21 RFU/min to 36 RFU/min and 34 RFU/min for 4% and 6% dextran, respectively. The synthesis rate dropped to 28 RFU/min and 25 RFU/min for 8% and 10% dextran. Ficoll had an optimal concentration at 6% and dextran in the range 4% - 6% in the PURE v3.50 formulation, indicating that we were operating at the optimal concentration.

We have shown that protein synthesis efficiency is increased in the diluted PURE systems (Fig 3g). The addition of crowding agents further increased these protein synthesis efficiencies drastically. The addition of dextran increased the efficiency from 0.48 to 0.71 (+48%) for PURE v3.90. Pure v3.50 protein synthesis efficiency increased from 0.34 to 0.86. This represents a 153% improvement in protein synthesis efficiency for the PURE v3.50 system with 6% dextran addition. This is particularly intriguing as the protein synthesis efficiency for PURE v3.50 without crowding agents was lower than PURE v3.90. It therefore appears that a high total non-ribosomal protein concentration of ∼200 µg/mL or above provided sufficient crowding, and that addition of crowding agents therefore had no effect, whereas the total non-ribosomal protein concentration in dilute PURE versions is not sufficient to provide sufficient crowding effects, and therefore addition of crowding agents enhanced synthesis rates in ultra-diluted PURE formulations.

Although we primarily focused on optimizing protein synthesis rates and protein synthesis efficiency in the context of self-regeneration and synthetic cell engineering, another parameter generally used for evaluating performance of cell-free systems is the total or final protein yield obtained in a batch reaction. We observed a considerable difference in protein yield for the various PURE formulations with and without addition of crowding agents (Fig 3h). In a concentrated PURE system such as PURE v1, the final yield without crowding agent was 115.2 µg/mL and dropped to 91.9 µg/mL (-20%) and 89.8 µg/mL (-22%), for 6% Ficoll and dextran, respectively. For PURE v3 without crowding agent, the final yield was 56 µg/mL. 6% dextran addition increased the final yield to 68.9 µg/mL (+23%). Ficoll addition did not have an impact on the final yield. In PURE v3.90 and v3.50, the final yields without crowding agent were 42.4 µg/mL and 22.8 µg/mL, respectively. Surprisingly, addition of dextran drastically increased these final yields to 68.4 µg/mL and 68.2 µg/mL. These correspond to 61% and 199% increases and super ceded the final yield of PURE v3 without addition of crowding agents. We thus were able to achieve high protein yields with PURE v3.50 which contains only 5.4% of the original PURE non-ribosomal proteins, but returned 59% of the total protein yield obtained with a regular PURE reaction (PURE v1).

Finally, we tested combinations of Ficoll and dextran to determine whether any synergistic effects on synthesis rate and yield could be achieved. Since 6% was found to be the optimal crowding agent concentration, we tested the following combinations of crowding agents: 3% Ficoll + 3% dextran, 6% Ficoll + 3% dextran, 3% Ficoll + 6% dextran, 6% Ficoll + 6% dextran. We tested these combinations in PURE v3.90 and PURE v3.50 and found that there was no additivity in synthesis rate or final yield (Supp Fig 14). We explored the combined effect of T7 RNAP concentration and dextran addition in v3.90 and v3.50 systems. New PURE systems were prepared with T7 RNAP at a concentration of 10 µg/mL (v3.90 T7 RNAP full conc and v3.50 T7 RNAP full conc) and tested with and without 6% dextran addition. As seen before, dextran addition in v3.90 with nominal T7 RNAP concentration increased synthesis rate from 10 RFU/min to 29 RFU/min. Similarly, in PURE v3.50 T7 RNAP full conc led to a slight increase in synthesis rate from 4 RFU/min to 8 RFU/min (Supp Fig 15). We also tested other potential crowding agents including 2,2,2-trifluoroethanol^37^ (TFE) at 1% - 6% (v/v) (Fig 3j), sucrose^38^ at 0.3 M, trimethylamine N-oxide^39^ (TMAO) and glycine^40^ at 100 mM and 1 M concentration in PURE v3.90 system (Fig 3i), but none of these crowding agents had a positive effect on synthesis rate or final yield.

### Self-regeneration capacity of the improved PURE systems

The main objective of this study was to optimize the protein synthesis efficiency of the PURE system by maximizing the ratio of protein synthesis rate to the total non-ribosomal protein content. We previously demonstrated self-regeneration of T7 RNA polymerase and the simultaneous self-regeneration of up to 7 essential AARSs on a microfluidic chemostat chip^9^. The synthesis capacity of PURE running in a chemostat microfluidic chip was determined to be 0.44 µg/mL/min with a doubling time of 47 minutes (exchanging 20% of the reactor every 15 minutes). Under these conditions, we estimated that 15 of 36 proteins could be self-regenerated in the PURE v1 system which accounts for 3.6% of the total PURE non-ribosomal protein concentration. These numbers slightly improved to 19 out of 36 proteins and 4.0% of total non-ribosomal protein content with the improved PURE v2 system (Fig 1e and Fig 4c).

**Figure 4:**
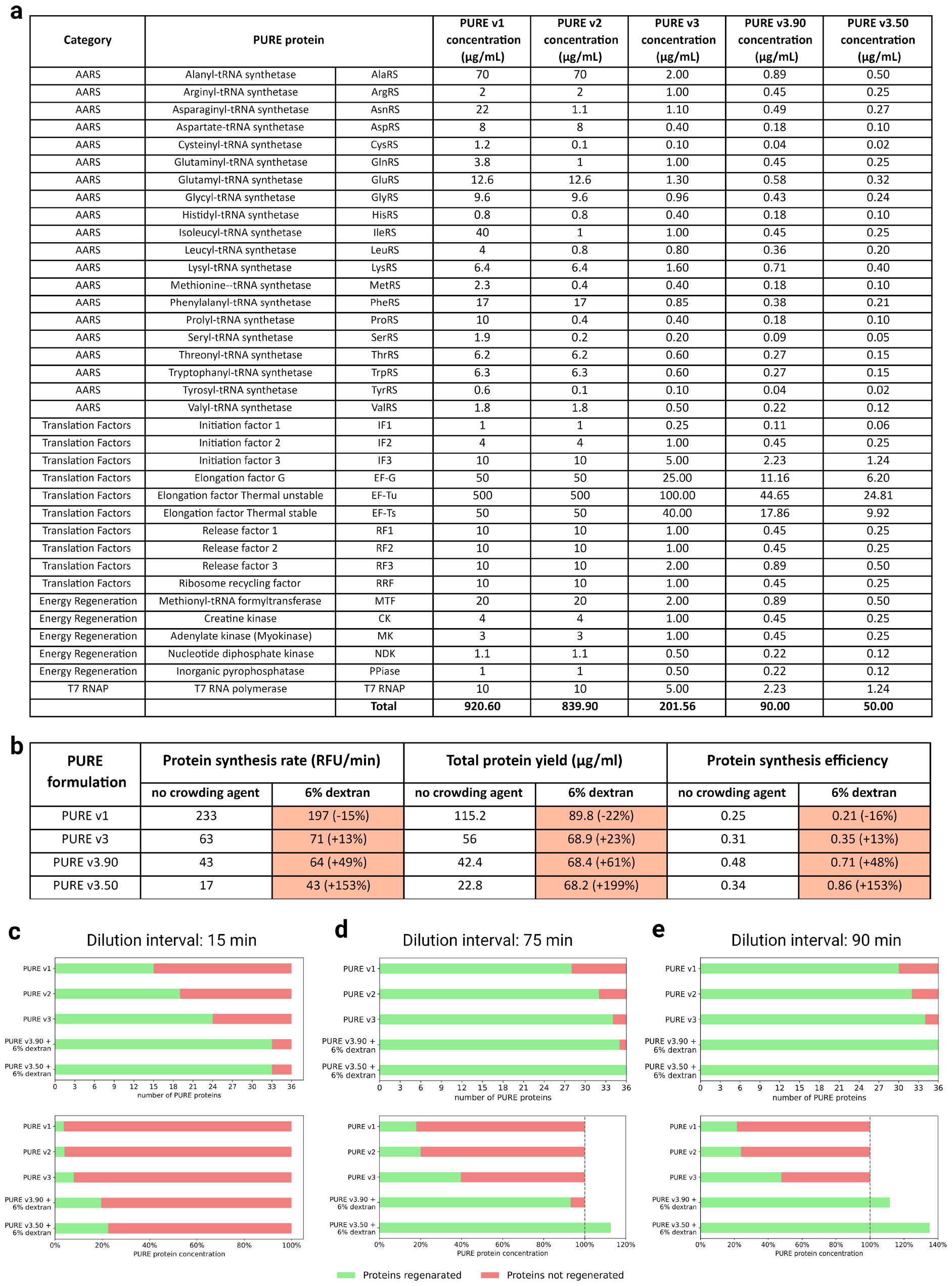
PURE system formulations and the self-regenerative capacity of selected PURE system formulations. (a) Table showing all 36 protein concentrations in the different PURE formulations. PURE v1 is the original formulation developed by Shimizu et al.^12^, PURE v2 a slightly optimized formulation previously published^9^, and PURE v3 and its dilution derivatives v3.90 and v3.50 were developed here. The total non-ribosomal protein concentration of PURE v3.90 and v3.50 were reduced by over one order of magnitude compared to PURE v1. (b) Table showing the performance summary of different PURE formulations with and without 6% dextran addition. (c-e) The theoretical self-regeneration capacity of different PURE formulations running in a chemostat with (c) 15-minute dilution intervals, (d) 75 min dilution interval, and (e) 90 min dilution interval. The graphs in the top row show the number of proteins regenerated and the bottom row graphs show the percentage of total non-ribosomal protein concentration that can be regenerated. PURE v3.90 and 3.50 are now theoretically capable of regenerating all of the PURE non-ribosomal protein content.

We evaluated protein self-regeneration in the PURE formulations developed here and listed in Fig 4a. The performance summary of PURE formulations including protein synthesis rate, total protein yield and protein synthesis efficiency with 6% dextran addition are listed in Fig 4b. For PURE v3, the synthesis rate was found to be roughly half when compared to PURE v1 under batch conditions. Assuming this relation holds for chemostat conditions as well, we estimated the chemostat synthesis rate of PURE v3 to be 0.213 µg/mL/min. Under these assumptions PURE v3 can regenerate 24 of 36 proteins, representing 7.9% of total PURE protein concentration. The PURE v3.90 system with 6% dextran crowding agent was estimated to have a chemostat synthesis rate of 0.223 µg/mL/min allowing 33 of 36 proteins to be regenerated representing 18.6% of the total PURE non-ribosomal protein concentration and the PURE v3.50 formulation with 6% dextran addition was estimated to have chemostat protein synthesis capacity of 0.150 µg/ml/min allowing 33 of 36 proteins to be regenerated or 22.5% of the total PURE non-ribosomal protein concentration (Figure 4c).

We recently demonstrated continuous exchange of small molecules by dialysis on a second-generation chemostat and measured PURE system performance on this device. A PURE reaction could be run on this device with 4 times longer dilution interval (dilutions every 60 minutes instead of every 15 minutes) giving rise to a doubling time of 186 minutes as opposed to 47 minutes while maintaining the same protein synthesis rate^25^. As synthesis rates remain constant but dilution rates are slowed by a factor of 4 it should now theoretically be possible to self-regenerate 35 non-ribosomal proteins when using PURE v3.90 and v3.50 system with 6% dextran addition, EF-Tu being the only exception. Furthermore, synthesis capacity is estimated to be 74.5% and 90.1% for v3.90 and v3.50 respectively. It should be theoretically possible to self-regenerate all 36 proteins in PURE v3.50 + 6% dextran when the dilution interval is further increased to 75 min (235 minute doubling time) leading to an estimated synthesis capacity of 112.6% (Fig 4d). With 90 min dilution intervals (282 minute doubling time), all 36 proteins can be theoretically self-regenerated in PURE v3.90 and v3.50 + 6% dextran and their synthesis capacity is estimated to be 111.7% and 135.1% respectively (Fig 4e). These doubling times are still well within the range of doubling times of naturally occurring bacteria (M. tuberculosis has a doubling time of about 1440 minutes for example). The synthesis capacity of diluted PURE systems v3.90 and v3.50 with dextran addition under longer dilution intervals thus theoretically exceeds the total non-ribosomal protein content of the PURE system with additional synthesis capacity for the implementation of genetic circuits for example.

## Discussion

Considerable progress is being made towards the bottom-up development of living systems. One of the core functions a living system has to fulfill is self-regeneration or autopoiesis^41^. A major milestone is therefore the development of a self-regenerating biochemical systems or universal biochemical constructor^1^. The PURE system is an ideal starting point for the development of a biochemical constructor as it is a defined system that consists of relatively few components. Considerable progress is being made in implementing DNA replication^5,6^, and tRNA synthesis^20^ in the PURE system, and some recent work has attempted to establish ribosome biogenesis *in vitro*^16,17,18,19^. We focused on self-regeneration of the non-ribosomal PURE proteins, and recently demonstrated that it is possible to robustly self-regenerate up to 7 essential AARSs in a microfluidic chemostat for extended periods of time^9^. But it also became clear that self-regeneration of all 36 non-ribosomal proteins of the PURE systems is a major challenge, mainly because the protein synthesis capacity of the PURE system is orders of magnitude too low compared to its own protein content. We therefore developed a novel microfluidic chemostat system that incorporates dialysis, allowing the PURE system to run under steady-state conditions at considerably lower dilution rates as previously achieved^25^. Lower dilution rates correspondingly lead to lower protein dilution rates, which in turn requires lower synthesis rates to achieve steady state levels.

In this work, we optimized the PURE system towards optimal self-regeneration. Unlike previous approaches of PURE system optimization, we didn’t focus on optimizing final protein yield, which was previously achieved by increasing the total non-ribosomal protein concentration of the system. We instead optimized the system’s protein synthesis efficiency which we define as the ratio of protein synthesis rate to total non-ribosomal protein concentration of the system.

We individually optimized all 36 PURE protein concentrations and arrived at a new PURE v3 with a total non-ribosomal protein content of 201.56 µg/mL, a reduction of 78% compared to the original PURE v1 formulation. Although we reduced total protein concentration almost five-fold the protein synthesis efficiency of PURE v3 is twice that of PURE v1. Further, we explored the limits of the PURE system. PURE v3 was diluted to 90 µg/mL (PURE v3.90), 50 µg/mL (PURE v3.50), 25 µg/mL (PURE v3.25) and, 10 µg/mL (PURE v3.10). We found that PURE remained functional at a concentration of 25 µg/mL. The synthesis rate of PURE v3.90 was around 60 RFU/min with the highest protein synthesis efficiency of 0.48 compared to the other PURE formulations. This protein synthesis efficiency is almost two-fold higher than PURE v1 (0.25).

We then explored the effect of crowding agents in these newly developed PURE formulations. Although crowding agents have not been particularly effective at improving PURE system performance in the past, we hypothesized that they could potentially be effective in these very dilute PURE formulations. Addition of crowding agents to PURE v1 or even PURE v3 did not lead to any enhancements, but when adding crowding agents to PURE v3.90 it resulted in an increase in protein synthesis rate. We tested the addition of both Ficoll and dextran, as well as combinations thereof, and found the optimal crowding agent to be dextran at a concentration of 6%. Dextran increased the protein synthesis rate of PURE v3.90 from 43 RFU/min to 64 RFU/min (+49%) and more than doubled the protein synthesis rate of PURE v3.50 from 17 RFU/min to 43 RFU/min (+153%). These correspond to increases in protein synthesis efficiency from 0.48 to 0.71 for PURE v3.90 and from 0.34 to 0.86 for PURE v3.50, making the latter the most efficient system to date. Total protein yield was also improved, with a 61% increase in total yield for PURE v3.90 (from 42.4 µg/mL to 68.4 µg/mL) and almost tripling (199%) the total yield of PURE v3.50 (from 22.8 µg/mL to 68.2 µg/mL). Both of these systems, despite being diluted by more than an order of magnitude compared to PURE v1, achieved similar total protein yields as the undiluted PURE v3, and approached the expression levels of the original PURE v1 formulation.

These new PURE formulations demonstrate that the PURE system is extremely robust to perturbations and modifications and make creating the PURE system even easier, allowing researchers to generate larger PURE volumes, particularly when combined with the OnePot method^13^. The new PURE formulations should therefore be useful for many applications in which maximal protein yield is not the primary concern, which encompasses the majority of PURE applications in synthetic biology. It is also intriguing that the PURE system sustains significant protein synthesis under very dilute conditions, which is of potential interest to researchers studying the origin of life where the earliest biochemical system was likely considerably more dilute than current cellular environments. Finally, all of the systems drastically outperform the original PURE formulation in terms of protein synthesis efficiency. Together with the recently developed dialysis based microfluidic chemostat, we estimate that it should now be theoretically possible to achieve full self-regeneration of all 36 non-ribosomal proteins of the PURE system on a microfluidic dialysis chemostat combined with longer dilution intervals which marks a major milestone towards the development of a living synthetic cell.

## Acknowledgements

We would like to thank Yoshihiro Shimizu (RIKEN) for kindly providing plasmids for the PURE system and Harris Wang (Wyss Institute, Harvard) for kindly providing *E. coli* RB1 strain for his-tag ribosome purification. We also thank Barbora Lavickova for providing some of the purified PURE protein components. The graphical schematic representations are created using BioRender.com. This work was supported by the European Research Council under the European Union’s Horizon 2020 research and innovation program grant 723106, a Swiss National Science Foundation Sinergia Grant (514558) and a Swiss National Science Foundation MINT grant (514337).

## Author contributions

R.B.G. performed experiments. R.B.G. and S.J.M. designed experiments, analyzed data, and wrote the manuscript.

## Methods

### Materials

For protein expression, *E. coli* strains BL21 and M15 were used. *E. coli* RB1 strain^42^ was used for his-tag ribosome purification and was obtained from Harris Wang (Wyss Institute, Harvard University, USA). Plasmids encoding 36 PURE proteins were obtained from Yoshihiro Shimizu (RIKEN Center for Biosystems Dynamics Research, Japan). For *in vitro* eGFP synthesis, a plasmid encoding the gene sequence was prepared by extension PCR from pKT127 plasmid as previously described^43^ and subsequently cloned into pSBlue-1 plasmid. This newly encoded eGFP plasmid was used for obtaining linear DNA templates by PCR amplification. The eGFP sequence and primer sequences used for PCR are provided in Supplementary Table 6. PCR amplified linear template DNA sequences were purified using DNA Clean and Concentrator-25 kit (Zymo Research). The resulting purified DNA was eluted in nuclease-free ultrapure water. DNA concentration was measured by UV-Vis absorbance at 260 nm using a Nanodrop instrument (ThermoFisher).

### His-tag ribosome purification

The purification procedure and buffers used in this work were prepared as previously described with slight modifications^9,44^. All buffers used are provided in Supplementary Table 4. β-mercaptoethanol was added just before use. Buffers were filtered with Flow Bottle Top Filters 0.45 µm aPES membrane and stored at 4°C. An overnight culture of *E. coli* RB1 strain was grown in 5 mL LB media at 37°C with shaking at 260 RPM. Purification culture (2 L) was prepared in 4 ^*^ 1 L baffled flask, each with 500 mL LB media and 3 mL overnight Rb1 culture. Culture was incubated at 37°C with shaking at 260 RPM and grown to exponential phase (3-4 hours). The cells were centrifuged at 3220 RCF for 20 min at 4°C and the cell pellet stored at -80°C. For cell lysis, the cell pellet was resuspended in 15 mL lysis buffer, and sonicated on ice, using a Vibra cell 75186 with a probe tip diameter of 6 mm at 70% amplitude for 11 cycles of 20 s pulse ON and 20 s pulse OFF. Lysed cells were centrifuged at 21130 RCF for 20 min at 4°C to remove cell debris resulting from the cell lysate. Nickel-NTA gravity-flow chromatography was used for purification. The ribosome purification column was prepared using 5 mL of IMAC Sepharose 6 Fast Flow beads (GE Heathcare) charged with 0.1 N Nickel sulphate solution. The column was equilibrated with 30 mL lysis buffer. Cell lysate was loaded onto the column and washed with 30 mL lysis buffer. For removing weakly bound molecules, buffers with increasing imidazole concentration were prepared by mixing lysis buffer and elution buffer as follows. The column was washed with 30 ml wash buffer 1 (29 mL lysis buffer + 1 mL elution buffer) (5 mM Imidazole), 60 mL wash buffer 2 (50 mL lysis buffer + 10 mL elution buffer) (25 mM Imidazole), 30 mL wash buffer 3 (22 mL lysis buffer + 8 mL elution buffer) (40 mM Imidazole), 30 mL wash buffer 4 (18 mL lysis buffer + 12 mL elution buffer) (60 mM Imidazole). The ribosomes were eluted with 7.5 ml elution buffer (150 mM Imidazole) and placed on ice at all times. The ribosomes were transferred to a 15 mL Amicon Ultra filter unit 3kDa MWCO (Sigma-Aldrich) and centrifuged at 3220 RCF, 45 min and 4°C. The eluted ribosome solution was concentrated to ∼1 mL. The ribosome solution was subjected to buffer exchange using ribosome buffer to lower the imidazole concentration. The concentrated ribosome fraction (∼1 mL) was diluted to 15 mL using ribosome buffer and centrifuged as mentioned above. This step was repeated thrice. The resulting ribosome solution (∼1 mL) was further concentrated using 0.5 mL Amicon Ultra Filter unit 3 KDa MWCO (Sigma-Aldrich) by centrifugation at 14,000 RCF and 4°C. Ribosome concentration was measured by absorbance at 260 nm of 1:100 diluted sample. An absorbance of 10 of the diluted sample corresponded to 23 µM in undiluted sample. The ribosome concentration was finally adjusted to 3.45 µM for *in vitro* protein expression. The final yield is usually around 750 µL.

### PURE protein purification

The purification procedure and buffers used in this work are prepared as previously described with slight modifications^9,44^. All buffers used are provided in Supplementary Table 4. β-mercaptoethanol was added just before use. Buffers were filtered with Flow Bottle Top Filters 0.45 µm aPES membrane and stored at 4°C. An overnight *E. coli* culture was prepared in 5 mL LB media supplemented with 100 µg/mL ampicillin and incubated at 37°C with shaking at 260 RPM. An *E. coli* purification culture (2 L) was prepared in 4 ^*^ 1 L baffled flasks, each with 500 mL LB media supplemented with 100 µg/mL ampicillin and 3.6 mL overnight culture. The culture was incubated at 37°C with shaking at 260 RPM for 2 hours and induced with 0.1 mM isopropyl β-D-1-thiogalactopyranoside (IPTG) for 3 hours. The culture was then centrifuged at 3220 RCF for 20 min at 4°C and the cell pellet stored at -80°C. The cell pellet was then resuspended in 15 mL Buffer A. Cells were lysed via sonication on ice using Vibra cell 75186 with probe tip diameter 6 mm at 70% amplitude for 11 cycles of 20 s ON pulse and 20 s OFF pulse. Cells were centrifuged at 25,000 RCF for 20 min at 4°C to remove cell debris. PURE protein was purified using Ni-NTA gravity flow chromatography. Purification column was prepared using 2-3 mL IMAC Sepharose 6 Fast Flow beads (GE Healthcare) and charged with 0.1 N Nickel sulphate solution. The column was then equilibrated with 30 mL buffer A solution. Supernatant from the cell lysate was loaded onto the column and the column washed with 25 mL buffer A solution, and 50 mL wash buffer 1 (47.5 mL buffer A + 2.5 mL buffer B). The protein was eluted with 10 mL elution buffer (1 mL buffer A + 9 mL buffer B). The eluted protein was subjected to buffer exchange by dialysis. A cellulose membrane with avg. flat width 33 mm with 14,000 MWCO (Merck) was used as dialysis tube by cutting two pieces of the dialysis membrane tubing with length ∼7 cm each. The tubing was soaked in HT buffer for 5 min. In each tube, one end was closed, filled with 5 mL eluted proteins, followed by closing of the other end. The dialysis tubes with protein were placed in a 1 L HT buffer overnight with shaking at 150 RPM and 4°C. The HT buffer was exchanged with 1 L stock buffer and placed for 3 hours under shaking at 150 RPM at 4°C. The protein concentration was measured by absorbance at 280 nm. In case higher concentrations were needed, the protein was concentrated using 0.5 mL Amicon Ultra Filter unit 3 KDa MWCO by centrifugation at 14,000 RCF and 4°C. The protein concentration used in different PURE versions are provided in Supplementary Table 1 (µg/mL) and Supplementary Table 2 (nM).

### Ni-NTA resin preparation, column regeneration and cleaning

For ribosome and PURE protein purification, column was prepared as follows. IMAC Sepharose 6 Fast Flow beads (GE Healthcare) was added into (Ribosome purification: 5 mL; PURE proteins purification: 2-3 mL) Econo-Pac chromatography columns (Biorad). The column was washed with 50 mL deionized Millipore water and charged with 15 mL 0.1 N Nickel sulphate solution. The charged column was washed with 50 mL deionized Millipore water to remove excess nickel sulphate. After purification, the column was regenerated as follows: washed with 30 mL deionized Millipore water, 10 mL buffer (0.2 M EDTA + 0.5 M NaCl), 30 mL 0.5 M NaCl solution, and 30 mL of deionized Millipore water. The column was stored in 20% (v/v) ethanol at 4°C. The column was cleaned as follows: wash with 30 mL deionized water, 30 mL 0.5 M NaOH solution, 30 mL deionized water, 30 mL 0.1 M acetic acid, 30 mL deionized water, 30 mL 70% (v/v) ethanol, 50 mL deionized water and store the column in 20% (v/v) ethanol at 4°C.

### Energy solution preparation

Energy solution was prepared as previously described^44^. The concentration and supplier information for all components are provided in Supplementary Table 5. Briefly, energy solution was prepared at 2.5x concentration containing 125 mM HEPES (4-(2-hydroxyethyl)-1-piperazineethanesulfonic acid), 250 mM potassium glutamate, 29.5 mM magnesium acetate, 50 mM creatine phosphate, 0.05 mM folinic acid, 5 mM spermidine, 0.75 mM of each amino acid, 5 mM ATP, 5 mM GTP, 2.5 mM CTP, 2.5 mM UTP and 130 UA260/mL of tRNA.

### Plate reader *in vitro* protein expression experiments

All plate reader experiments were setup with a total volume of 10 or 20 µL at 34°C. PURE protein titration experiments and PURE subset swap experiments were conducted with 10 µL reaction volumes. A 10 µL PURE reaction was set-up using 1.8 µL of 3.45 µM his-tag ribosome (final concentration: 0.6 µM), 1.3 µL of PURE proteins (PURE v1, v2 and v3), 4 µL of 2.5x energy solution, eGFP DNA linear template (final concentration: 4 nM) and brought to a final volume of 10 µL with addition of water. PURE dilution and crowding agent experiments were conducted using commercial NEB *E. coli* ribosomes (P0763S) and 20 µL reaction volumes. For a 20 µL PURE reaction, 0.9 µL of NEB ribosomes at 13.3 µM (final concentration: 0.6 µM), 2.6 µL of PURE proteins (PURE v1, v2, v3), 8 µL of 2.5x energy solution, eGFP DNA linear template (final concentration: 4 nM) and brought to a final volume of 20 µL with addition of water. In case of PURE v3.90, v3.50, v3.25, and v3.10: 1.17 µL, 0.65 µL, 0.325 µL and 0.13 µL were added respectively for PURE proteins in a 20 µL reaction. Experiment duration was determined by reaching the plateau stage which typically occurred after 3 hours for titration experiments using PURE v1 and v2. For PURE v3, v3.90, v3.50, v3.25, and v3.10 experiment duration was anywhere from 10 to 16 hours. eGFP was measured (excitation: 485 nm, emission: 515 nm) on a SynergyMX platereader (BioTek). Synthesis rate was calculated by measuring maximum slope in a moving average period of 20 minutes. For final yield, a calibration curve was used to calculate total concentration of eGFP produced (Supp Fig 16). PURE versions with T7 RNAP at nominal concentration (10 µg/mL) are provided in Supplementary Table 3. For crowding agent addition in PURE, dextran-70 (Carl ROTH) and FicollRTM, Type 70 (Santa Cruz Biotechnology Inc.) were used to prepare 10%, 20%, 30%, 40% and 50% (w/v) in nuclease-free water. For a 20 µL PURE reaction, 4 µL crowding agent was added to the reaction mix to achieve the final concentration of 2% - 10%. For combinations of dextran and Ficoll experiments, 4 µL of each crowding agent was added to the reaction mix. Trimethylamine N-oxide (TMAO) (Merck) and glycine (Carl ROTH), were used to prepare 0.1 M and 1 M solution. For 2,2,2-Trifluoroethanol (TFE) (Sigma-Aldrich) 5%, 10%, 15%, 20%, 25%, 30% (v/v) in nuclease-free water were prepared. Sucrose (Sigma-Aldrich) was used to prepare 0.3 M solution. For experiments with crowding agents TMAO, glycine, TFE, and sucrose, 4 µL was added to the 20 µL PURE reaction mix.

