## Supplementary Information for "Towards self-regeneration: exploring the limits of protein synthesis in the PURE cell-free transcription-translation system"

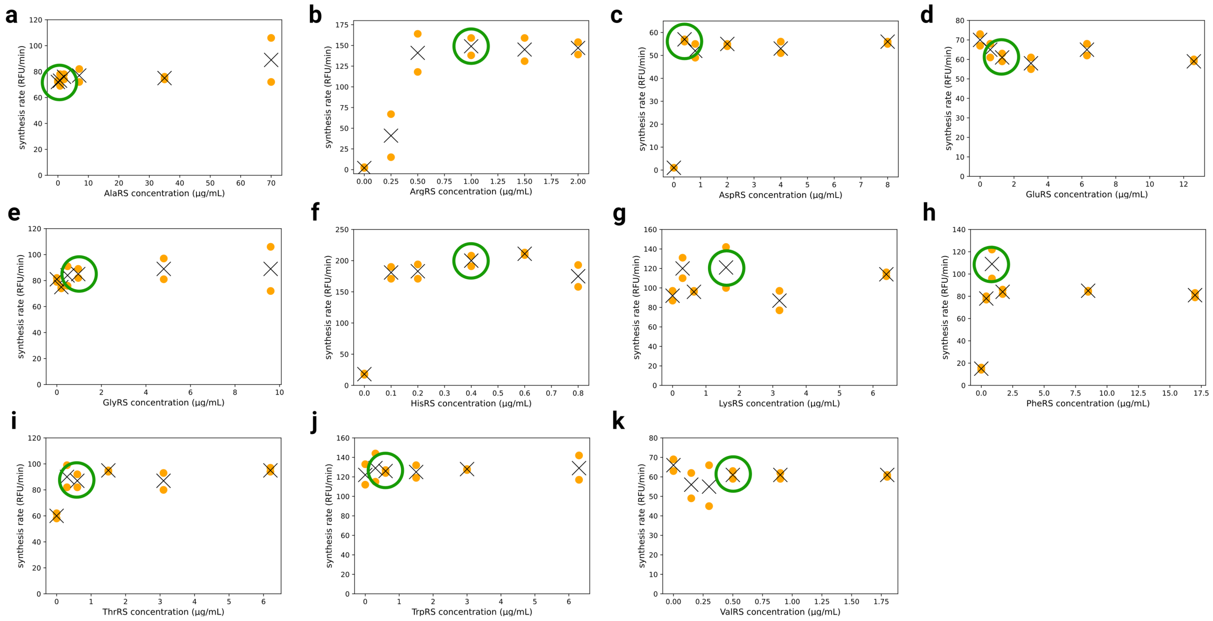


Supplementary Figure 1: PURE aminoacyl-tRNA-synthetase (AARS) titration plots. (a)-(k) Each of the 11 AARS proteins was titrated individually (the remaining 9 AARSs were previously optimized for PUREv2). An orange dot represents synthesis rate data points, black crosses represent the average synthesis rate and a green circle shows the chosen protein concentration for that particular AARS.

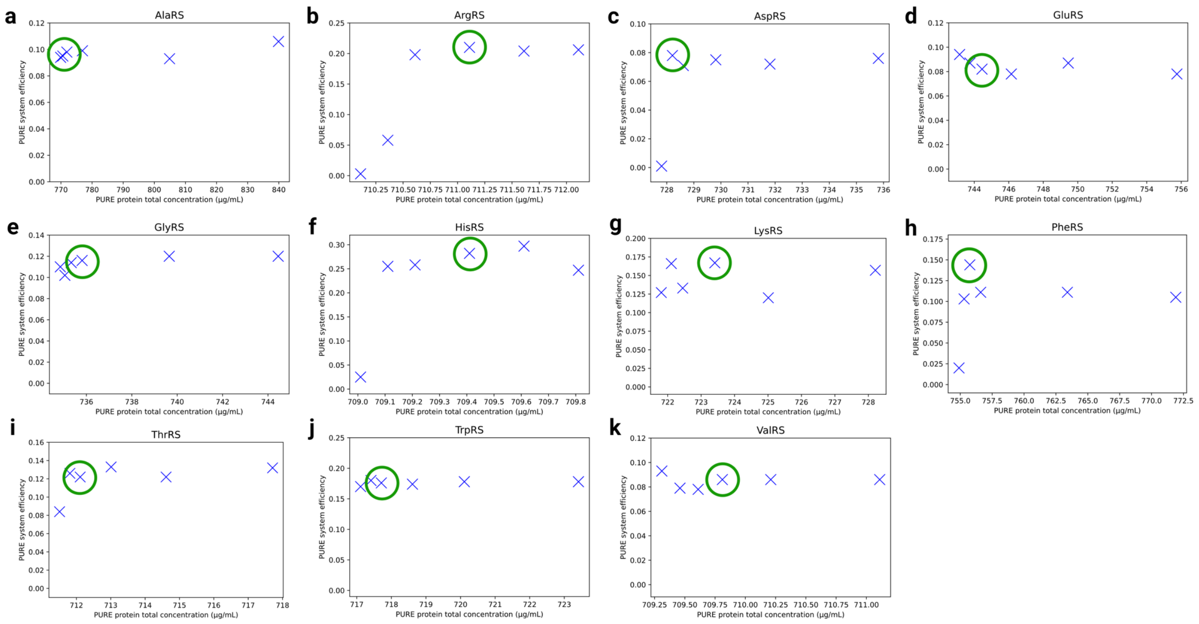


Supplementary Figure 2: Aminoacyl-tRNA-synthetase (AARS) efficiency plots. (a)-(k) For each of the 11 AARS proteins, protein synthesis efficiency was calculated as the ratio of average synthesis rate to total protein concentration. The green circles indicate the chosen protein concentration for a particular protein.

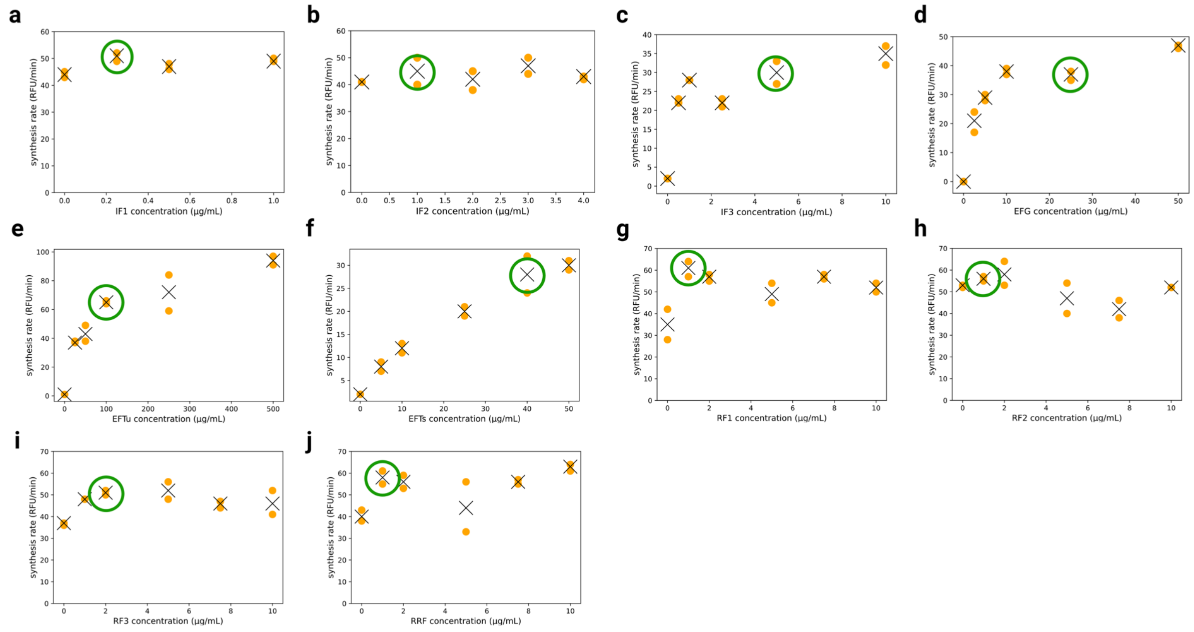


Supplementary Figure 3: PURE translation protein titrations. (a)-(j) Each of the 10 translation proteins in the PURE system was titrated individually. An orange dot represents synthesis rate data points, black crosses represent the average synthesis rate and a green circle shows the chosen protein concentration for that particular protein.

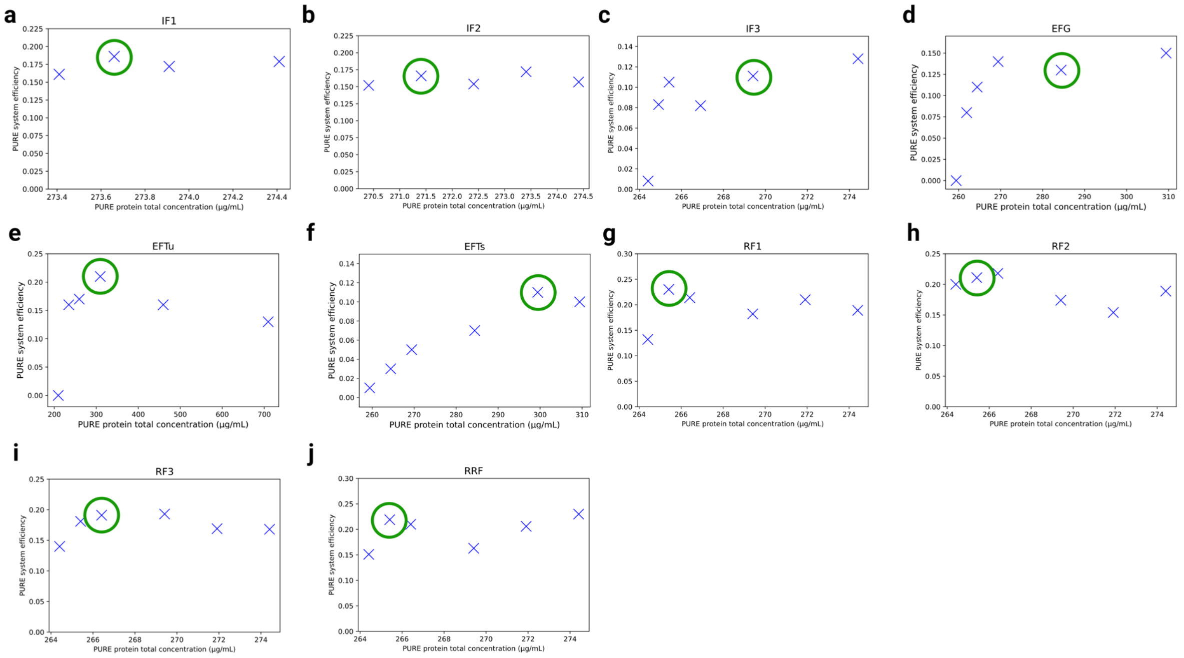


Supplementary Figure 4: PURE translation proteins efficiency plots. (a)-(j) For each of the 10 translation proteins, PURE system efficiency was calculated as the ratio of average synthesis rate to protein concentration titrated. The green circles indicate the chosen protein concentration for a particular protein.

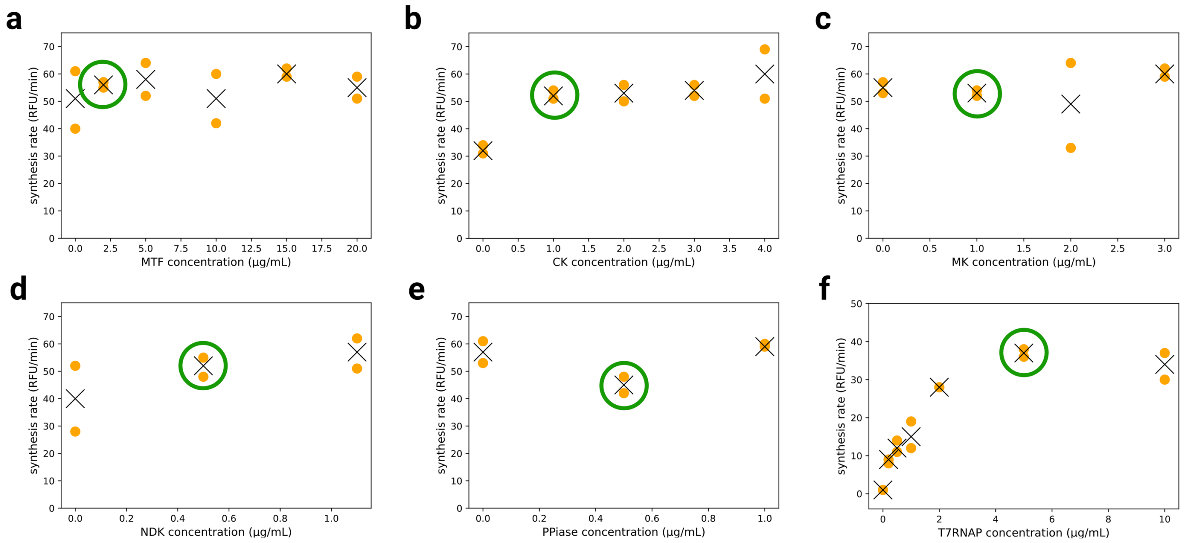


Supplementary Figure 5: PURE energy regeneration proteins and T7 RNA Polymerase titration plots. (a)-(e) For each of the 5 energy regeneration proteins, and, (f) T7 RNA Polymerase in the PURE system were titrated individually. An orange dot represents synthesis rate, black crosses represent the average synthesis rate and the green circles indicate the chosen protein concentration for a particular protein.

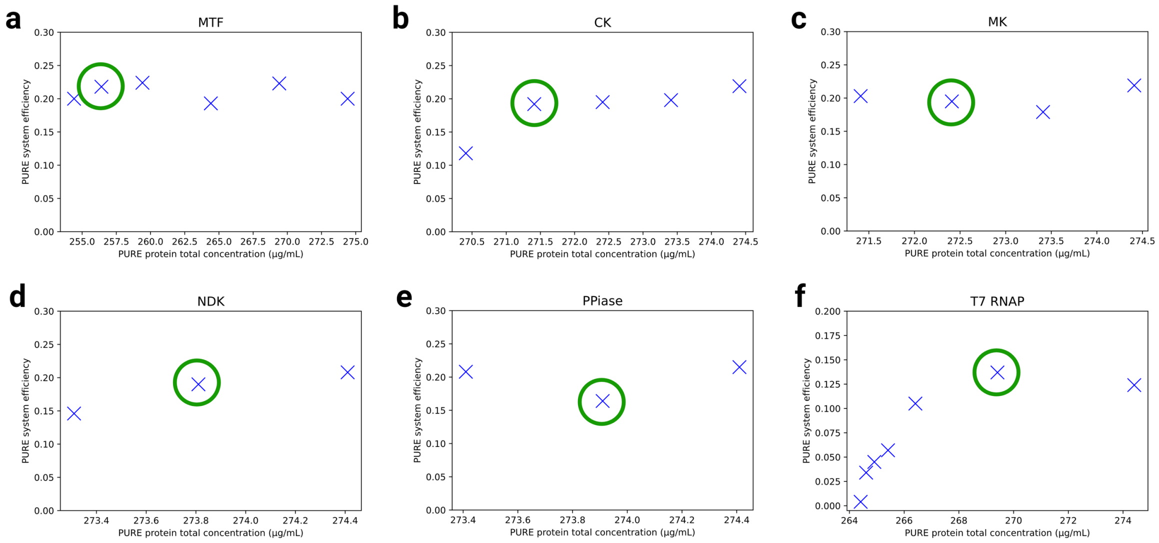


Supplementary Figure 6: PURE energy regeneration proteins and T7 RNA Polymerase efficiency plots. (a)-(e) Each of the 5 translation proteins, and, (f) T7 RNA Polymerase PURE system efficiency was calculated as the ratio of average synthesis rate to protein concentration. The blue crosses represent the PURE system efficiency plotted against total PURE protein concentration and the green circles indicate the chosen protein concentration for a particular protein.

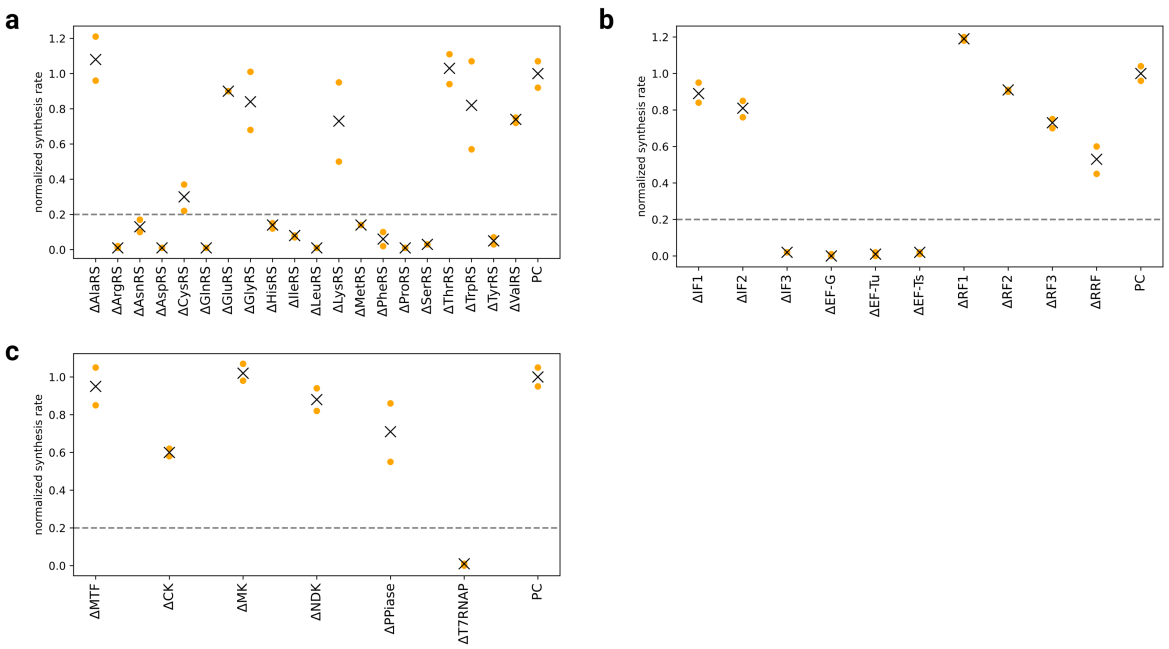


Supplementary Figure 7: PURE proteins dropout experiments. All 36 PURE proteins were individually omitted to check for essentiality. The synthesis rate was normalized with regard to the positive control. (a) aminoacyl-tRNA-synthetase (AARS), (b) translation factor proteins, (c) energy regeneration proteins and T7 RNA Polymerase. An orange dot represents synthesis rate, and black crosses represent the average synthesis rate. A dashed line shows the 20% cutoff we chose for considering whether a protein is essential or not. All proteins with an average synthesis rate below the dashed line were considered to be essential for protein expression.

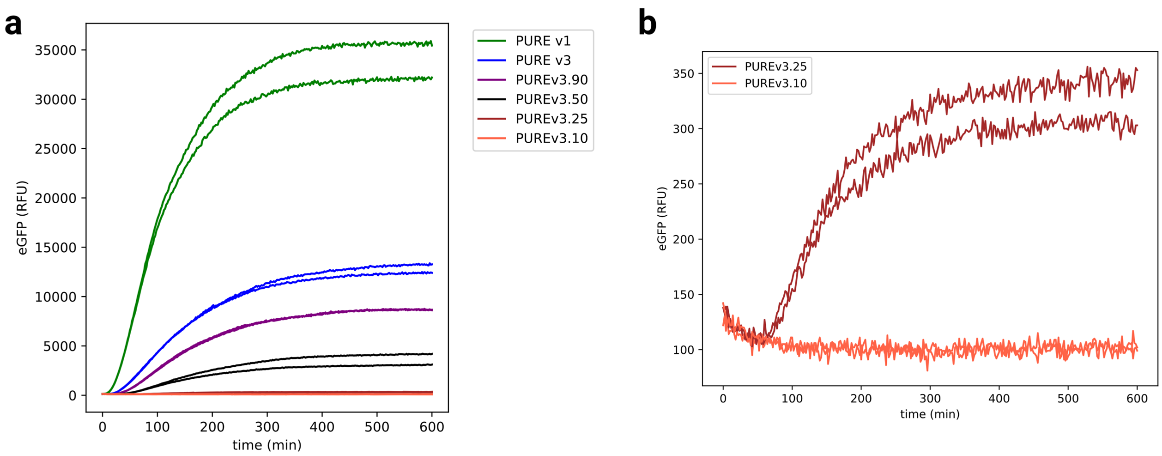


Supplementary Figure 8: PURE *in vitro* eGFP expression of different PURE formulations. (a) Plate reader eGFP fluorescent data for all PURE systems from PURE v1 to PURE v3.10. (b) eGFP expression data for PURE v3.25 and PURE v3.10. The experiment was performed in duplicates for each sample.

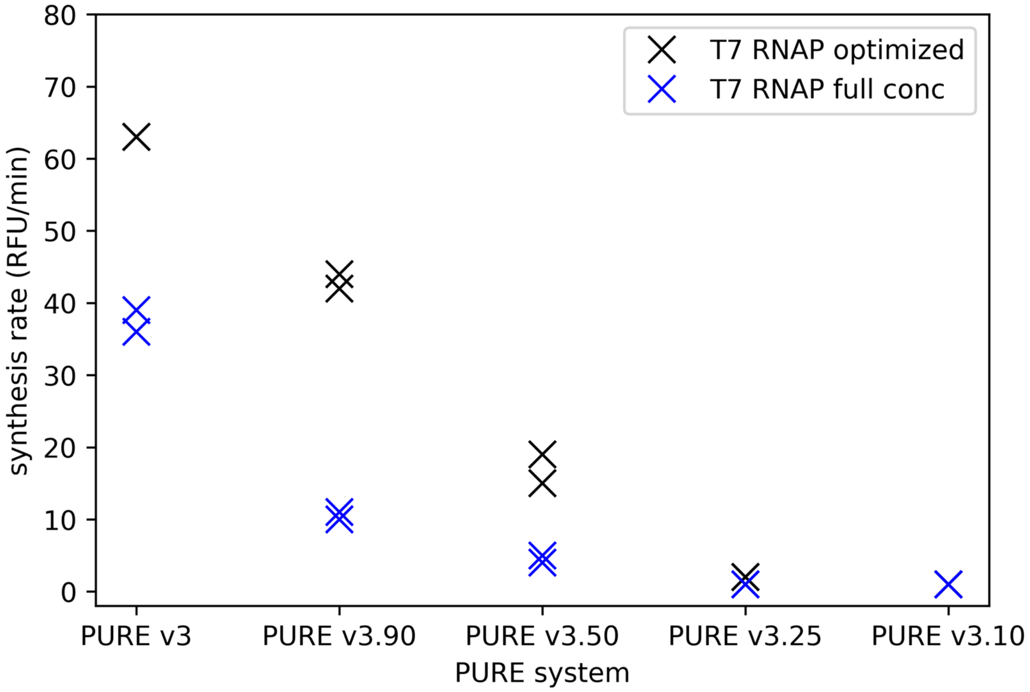


Supplementary Figure 9: PURE system synthesis rate with optimized T7 RNAP concentration (black) and full T7 RNAP concentration (blue). Optimized T7 RNAP concentration for different PURE formulation: 5 µg/mL (PURE v3), 2.23 µg/mL (PURE v3.90), 1.24 µg/mL (PURE v3.50), 0.62 µg/mL (PURE v3.25) and 0.25 µg/mL (PURE v3.10). Full T7 RNAP concentration for all PURE formulation is at 10 µg/mL. The experiment was performed in duplicates for each sample.

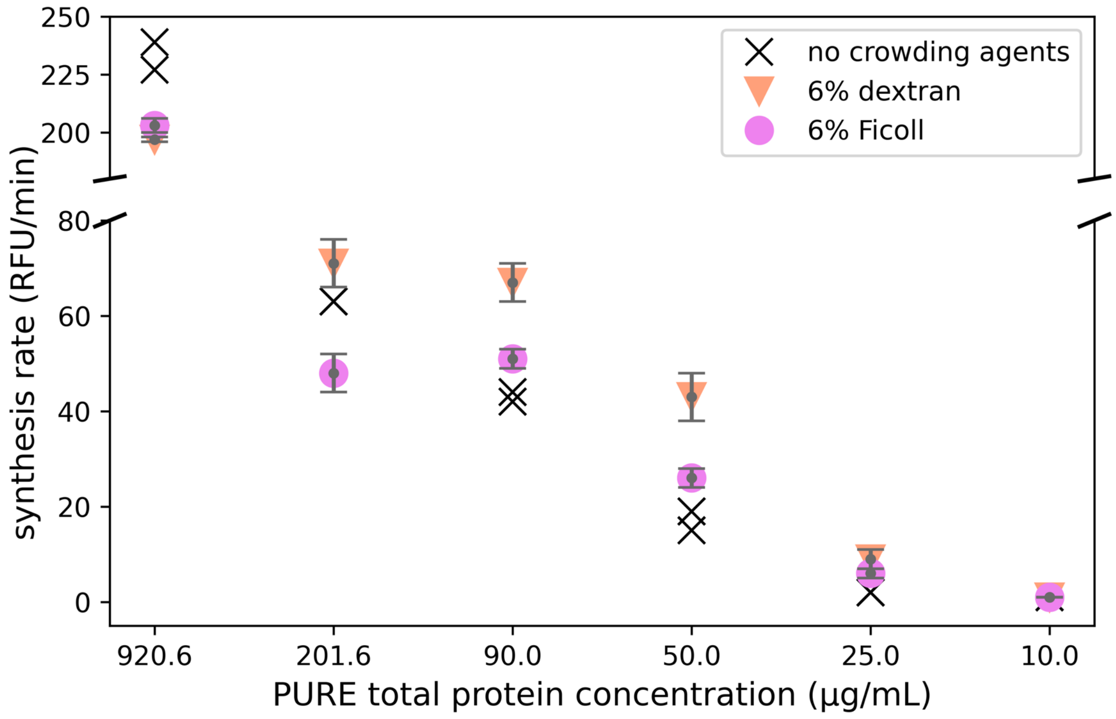


Supplementary Figure 10: PURE synthesis rate with crowding agents 6% dextran and 6% Ficoll added to different PURE formulations inlcuding PURE v1 with a reaction volume of 20 μL (n=2 for PURE without crowding agents and PURE v1 + 6% dextran group, n=4 for PURE with crowding agents except PURE v3.90 + 6% dextran, and n=10 for PURE v3.90 + 6% dextran group). Error bar presents mean ± standard deviation.

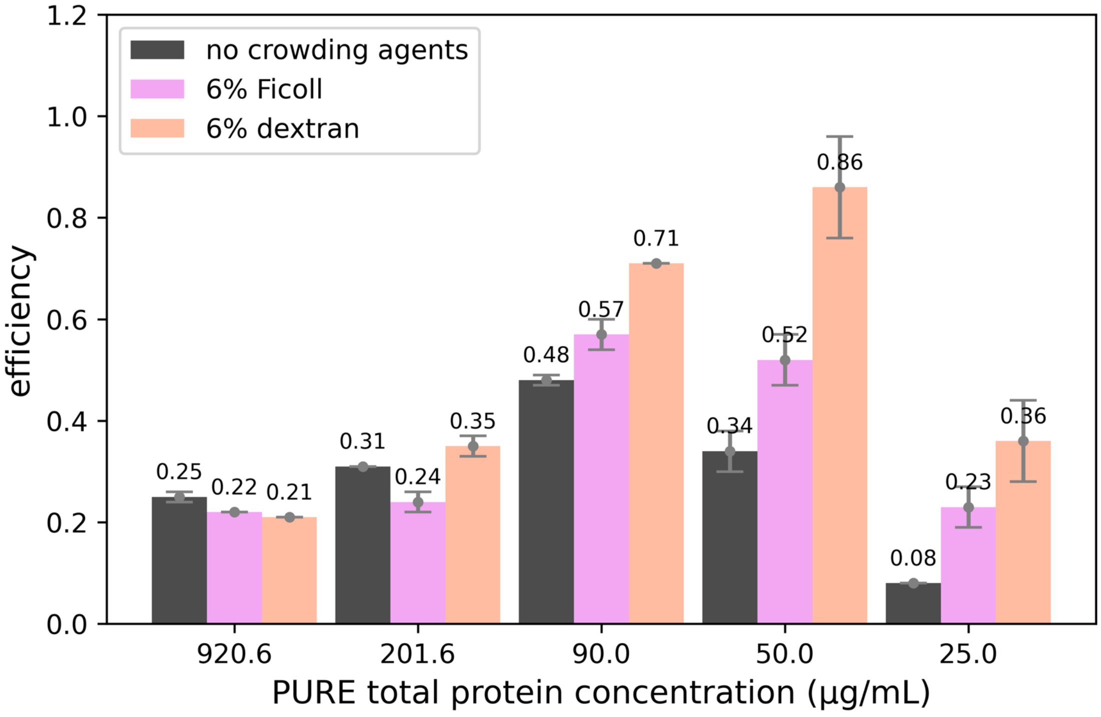


Supplementary Figure 11: PURE system efficiency with crowding agents 6% dextran and 6% Ficoll added to different PURE formulations inlcuding PURE v1 (n=2 for PURE without crowding agents and PURE v1 + 6% dextran group, n=4 for PURE with crowding agents except PURE v3.90 + 6% dextran, and n=10 for PURE v3.90 + 6% dextran group). Error bar presents mean ± standard deviation.

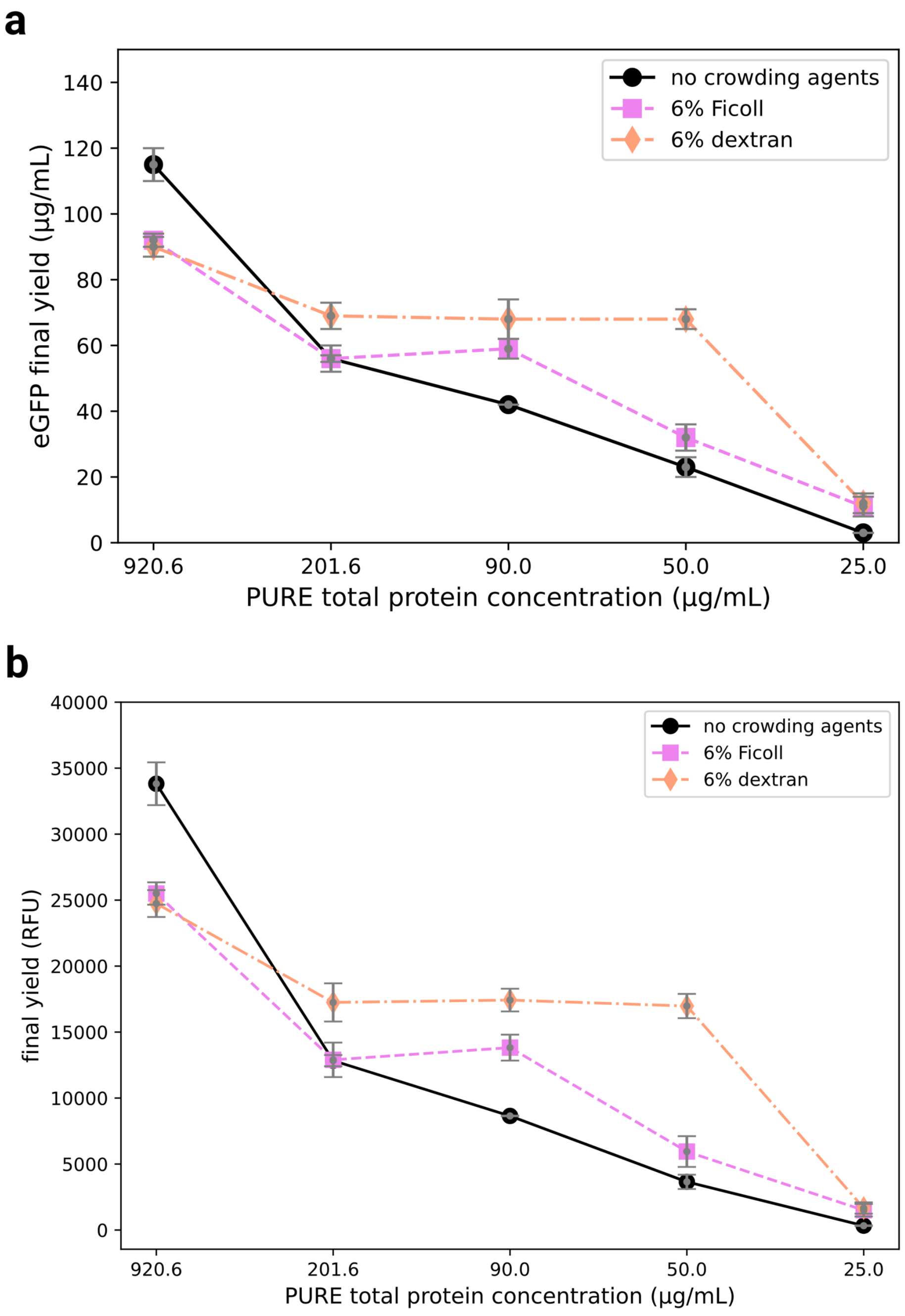


Supplementary Figure 12: PURE system eGFP final yield in µg/mL (a) and RFU (b) with and without 6% dextran and 6% Ficoll addition to different PURE formulations inlcuding PURE v1 with a reaction volume of 20 µL (n=2 for PURE without crowding agents and PURE v1 + 6% dextran group, n=4 for PURE with crowding agents except PURE v3.90 + 6% dextran, and n=10 for PURE v3.90 + 6% dextran group). Error bar presents mean ± standard deviation.

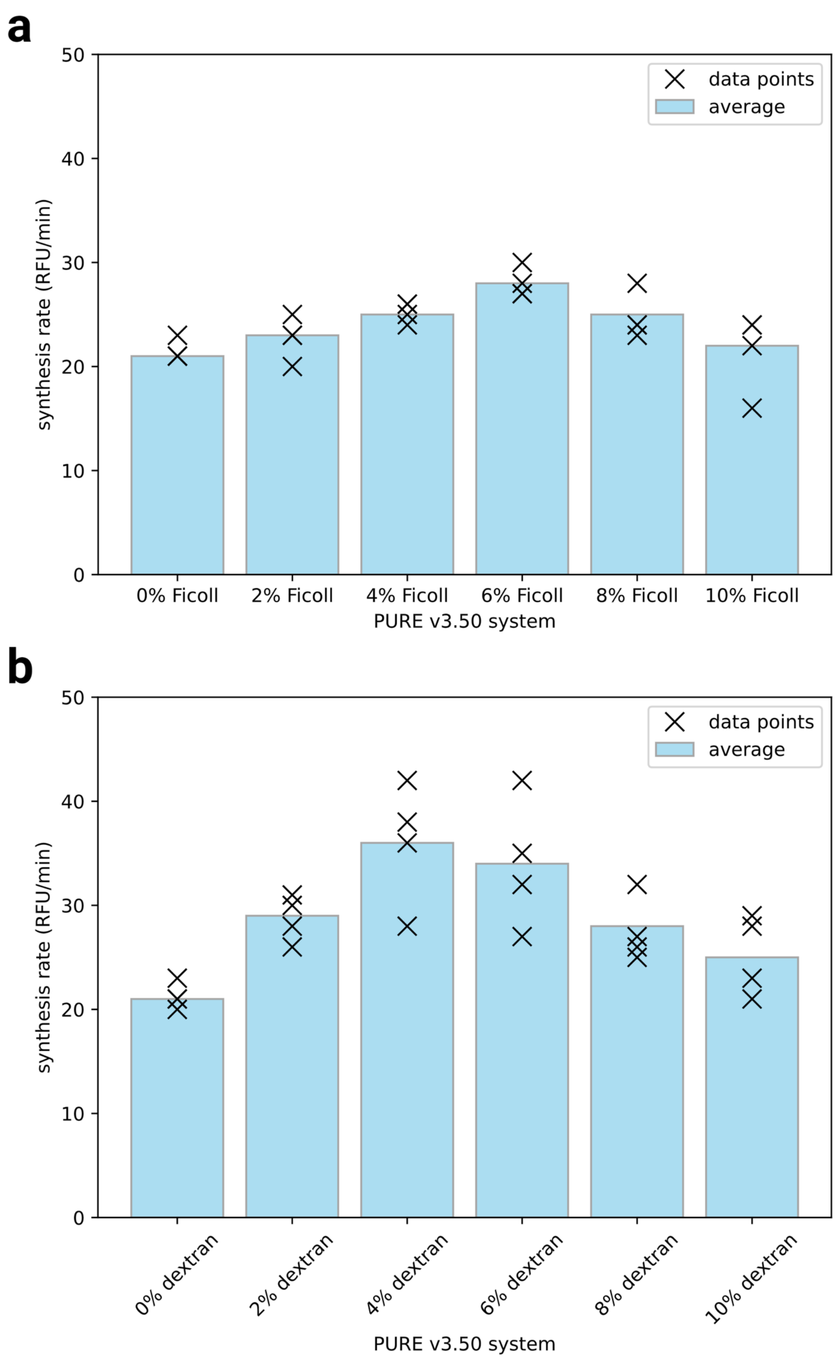


Supplementary Figure 13: Ficoll-70 (a) and dextran-70 (b) titration in the PURE v3.50 formulation with a reaction volume of 20 µL (n=4).

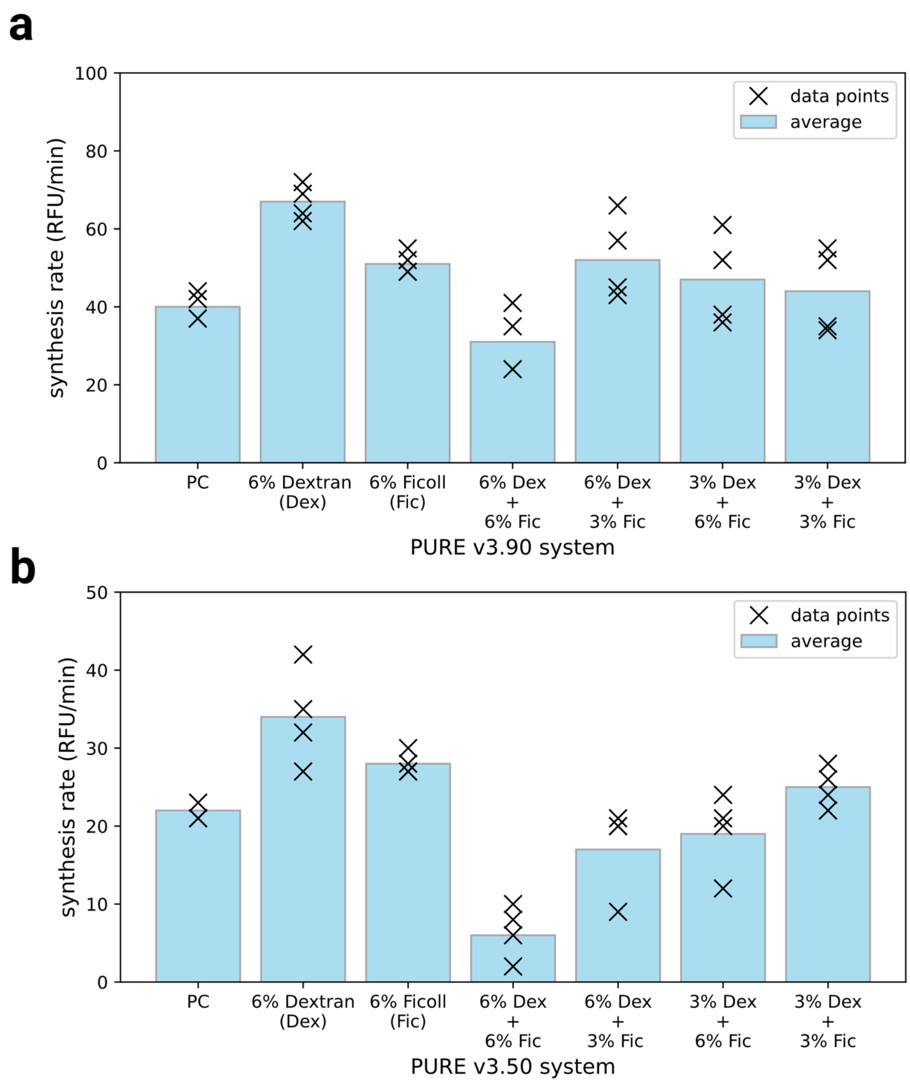


Supplementary Figure 14: Ficoll (Fic) and dextran (Dex) combination matrix in diluted (a) PURE v3.90 and (b) v3.50 with a reaction volume of 20 µL (n=4). The following combinations were tested on both systems (6% Dex, 6% Fic), (6% Dex, 3% Fic), (3% Dex, 6% Fic), (3% Dex, 3% Fic). The synthesis rate of 6% dextran, 6% Ficoll, and positive control (PC) are provided for reference. A black cross represents synthesis rate data points and a blue bar shows the average synthesis rate.

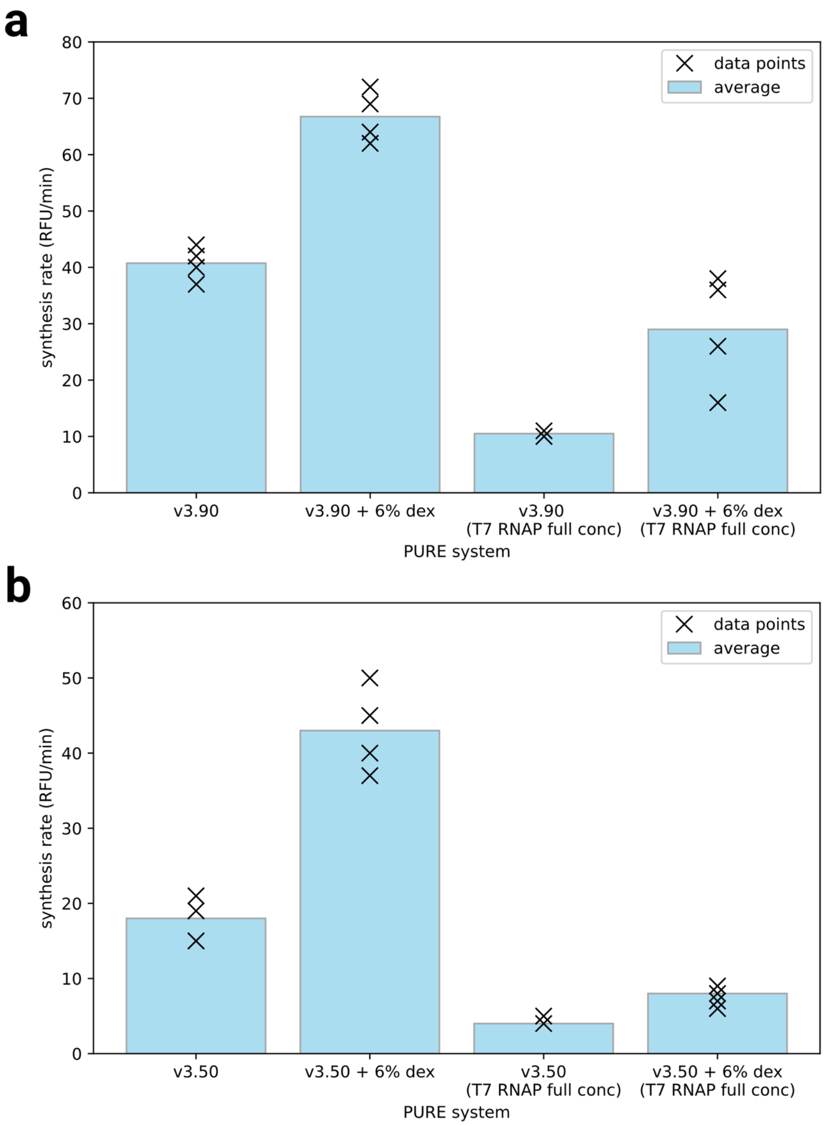


Supplementary Figure 15: Effect of T7 RNAP concentration on diluted PURE (a) v3.90 and (b) v3.50 with 6% dextran addition with a reaction volume of 20 µL. A higher concentration of T7 RNAP at 10 µg/ml (T7 RNAP full conc) did not have a positive impact compared to the optimal T7 RNAP concentration used in each diluted system. Dextran addition increased synthesis rate in both systems compared their counterparts without addition. T7 RNAP concentration in PURE v3.90 and PURE v3.50 is 2.23 µg/mL and 1.24 µg/mL respectively.

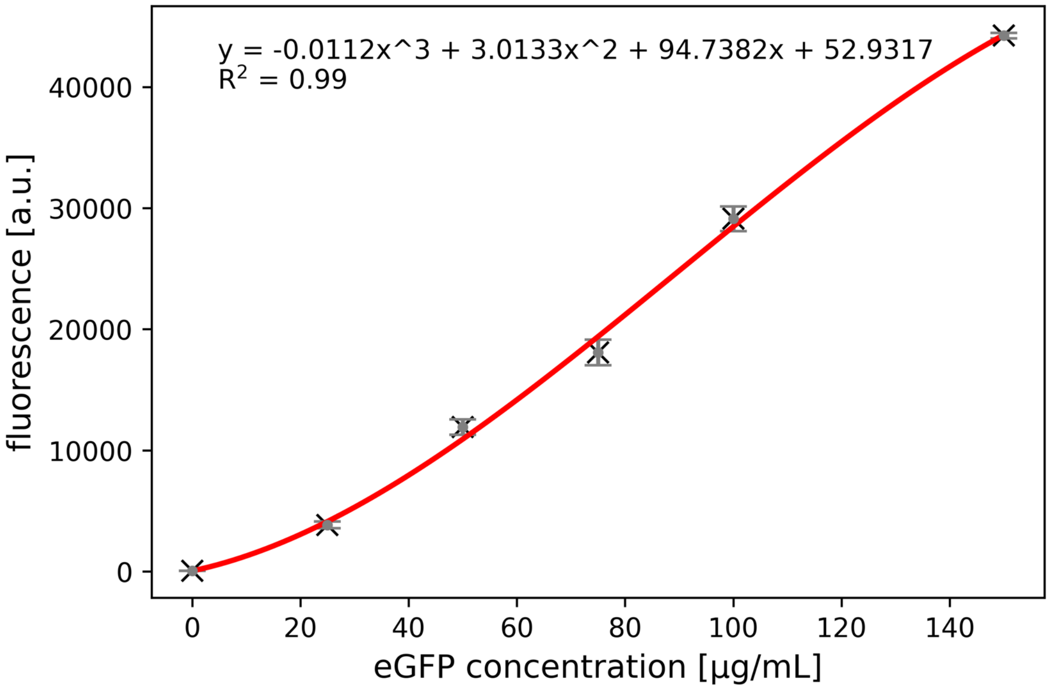


Supplementary Figure 16: Calibration curve for eGFP. The standard curve was produced by fluorescence measurement for eGFP (abcam 134853) in PBS buffer using a plate reader with the same settings as the *in vitro* protein expression experiment. Excitation and emission wavelengths were 485 and 515 nm respectively. Experiment were performed in triplicates. Data is shown as mean ± s.d. (n=3).

Supplementary Table 1: PURE protein concentrations for different PURE formulations in µg/mL.

| Category | PURE protein | | PURE v1 concentration (µg/mL) | PURE v2 concentration (µg/mL) | PURE v3 concentration (µg/mL) | PURE v3.90 concentration (µg/mL) | PURE v3.50 concentration (µg/mL) | PURE v3.25 concentration (µg/mL) | PURE v3.10 concentration (µg/mL) |
|---|---|---|---|---|---|---|---|---|---|
| AARS | Alanyl-tRNA synthetase | AlaRS | 70 | 70 | 2.00 | 0.89 | 0.50 | 0.25 | 0.10 |
| AARS | Arginyl-tRNA synthetase | ArgRS | 2 | 2 | 1.00 | 0.45 | 0.25 | 0.12 | 0.05 |
| AARS | Asparaginyl-tRNA synthetase | AsnRS | 22 | 1.1 | 1.10 | 0.49 | 0.27 | 0.14 | 0.05 |
| AARS | Aspartate-tRNA synthetase | AspRS | 8 | 8 | 0.40 | 0.18 | 0.10 | 0.05 | 0.02 |
| AARS | Cysteinyl-tRNA synthetase | CysRS | 1.2 | 0.1 | 0.10 | 0.04 | 0.02 | 0.01 | 0.00 |
| AARS | Glutaminyl-tRNA synthetase | GlnRS | 3.8 | 1 | 1.00 | 0.45 | 0.25 | 0.12 | 0.05 |
| AARS | Glutamyl-tRNA synthetase | GluRS | 12.6 | 12.6 | 1.30 | 0.58 | 0.32 | 0.16 | 0.06 |
| AARS | Glycyl-tRNA synthetase | GlyRS | 9.6 | 9.6 | 0.96 | 0.43 | 0.24 | 0.12 | 0.05 |
| AARS | Histidyl-tRNA synthetase | HisRS | 0.8 | 0.8 | 0.40 | 0.18 | 0.10 | 0.05 | 0.02 |
| AARS | Isoleucyl-tRNA synthetase | IleRS | 40 | 1 | 1.00 | 0.45 | 0.25 | 0.12 | 0.05 |
| AARS | Leucyl-tRNA synthetase | LeuRS | 4 | 0.8 | 0.80 | 0.36 | 0.20 | 0.10 | 0.04 |
| AARS | Lysyl-tRNA synthetase | LysRS | 6.4 | 6.4 | 1.60 | 0.71 | 0.40 | 0.20 | 0.08 |
| AARS | Methionine--tRNA synthetase | MetRS | 2.3 | 0.4 | 0.40 | 0.18 | 0.10 | 0.05 | 0.02 |
| AARS | Phenylalanyl-tRNA synthetase | PheRS | 17 | 17 | 0.85 | 0.38 | 0.21 | 0.11 | 0.04 |
| AARS | Prolyl-tRNA synthetase | ProRS | 10 | 0.4 | 0.40 | 0.18 | 0.10 | 0.05 | 0.02 |
| AARS | Seryl-tRNA synthetase | SerRS | 1.9 | 0.2 | 0.20 | 0.09 | 0.05 | 0.02 | 0.01 |
| AARS | Threonyl-tRNA synthetase | ThrRS | 6.2 | 6.2 | 0.60 | 0.27 | 0.15 | 0.07 | 0.03 |
| AARS | Tryptophanyl-tRNA synthetase | TrpRS | 6.3 | 6.3 | 0.60 | 0.27 | 0.15 | 0.07 | 0.03 |
| AARS | Tyrosyl-tRNA synthetase | TyrRS | 0.6 | 0.1 | 0.10 | 0.04 | 0.02 | 0.01 | 0.00 |
| AARS | Valyl-tRNA synthetase | ValRS | 1.8 | 1.8 | 0.50 | 0.22 | 0.12 | 0.06 | 0.02 |
| Translation Factors | Initiation factor 1 | IF1 | 1 | 1 | 0.25 | 0.11 | 0.06 | 0.03 | 0.01 |
| Translation Factors | Initiation factor 2 | IF2 | 4 | 4 | 1.00 | 0.45 | 0.25 | 0.12 | 0.05 |
| Translation Factors | Initiation factor 3 | IF3 | 10 | 10 | 5.00 | 2.23 | 1.24 | 0.62 | 0.25 |
| Translation Factors | Elongation factor G | EF-G | 50 | 50 | 25.00 | 11.16 | 6.20 | 3.10 | 1.24 |
| Translation Factors | Elongation factor Thermal unstable | EF-Tu | 500 | 500 | 100.00 | 44.65 | 24.81 | 12.40 | 4.96 |
| Translation Factors | Elongation factor Thermal stable | EF-Ts | 50 | 50 | 40.00 | 17.86 | 9.92 | 4.96 | 1.98 |
| Translation Factors | Release factor 1 | RF1 | 10 | 10 | 1.00 | 0.45 | 0.25 | 0.12 | 0.05 |
| Translation Factors | Release factor 2 | RF2 | 10 | 10 | 1.00 | 0.45 | 0.25 | 0.12 | 0.05 |
| Translation Factors | Release factor 3 | RF3 | 10 | 10 | 2.00 | 0.89 | 0.50 | 0.25 | 0.10 |
| Translation Factors | Ribosome recycling factor | RRF | 10 | 10 | 1.00 | 0.45 | 0.25 | 0.12 | 0.05 |
| Energy Regeneration | Methionyl-tRNA formyltransferase | MTF | 20 | 20 | 2.00 | 0.89 | 0.50 | 0.25 | 0.10 |
| Energy Regeneration | Creatine kinase | CK | 4 | 4 | 1.00 | 0.45 | 0.25 | 0.12 | 0.05 |
| Energy Regeneration | Adenylate kinase (Myokinase) | MK | 3 | 3 | 1.00 | 0.45 | 0.25 | 0.12 | 0.05 |
| Energy Regeneration | Nucleotide diphosphate kinase | NDK | 1.1 | 1.1 | 0.50 | 0.22 | 0.12 | 0.06 | 0.02 |
| Energy Regeneration | Inorganic pyrophosphatase | PPiase | 1 | 1 | 0.50 | 0.22 | 0.12 | 0.06 | 0.02 |
| T7 RNAP | T7 RNA polymerase | T7 RNAP | 10 | 10 | 5.00 | 2.23 | 1.24 | 0.62 | 0.25 |
| | | **Total** | **920.60** | **839.90** | **201.56** | **90.00** | **50.00** | **25.00** | **10.00** |

Supplementary Table 2: PURE protein concentrations for different PURE formulations in molar concentration. For PURE v3.50, v3.25, and v3.10 concentrations are provided in nM.

| Category | PURE protein | | PURE v1 concentration (μM) | PURE v2 concentration (μM) | PURE v3 concentration (μM) | PURE v3.90 concentration (μM) | PURE v3.50 concentration (nM) | PURE v3.25 concentration (nM) | PURE v3.10 concentration (nM) |
|---|---|---|---|---|---|---|---|---|---|
| AARS | Alanyl-tRNA synthetase | AlaRS | 0.714 | 0.714 | 0.020 | 0.009 | 5.06 | 2.53 | 1.01 |
| AARS | Arginyl-tRNA synthetase | ArgRS | 0.030 | 0.030 | 0.015 | 0.007 | 3.69 | 1.85 | 0.74 |
| AARS | Asparaginyl-tRNA synthetase | AsnRS | 0.408 | 0.020 | 0.020 | 0.009 | 5.06 | 2.53 | 1.01 |
| AARS | Aspartate-tRNA synthetase | AspRS | 0.119 | 0.119 | 0.006 | 0.003 | 1.48 | 0.74 | 0.30 |
| AARS | Cysteinyl-tRNA synthetase | CysRS | 0.023 | 0.002 | 0.002 | 0.001 | 0.47 | 0.23 | 0.09 |
| AARS | Glutaminyl-tRNA synthetase | GlnRS | 0.059 | 0.015 | 0.015 | 0.007 | 3.84 | 1.92 | 0.77 |
| AARS | Glutamyl-tRNA synthetase | GluRS | 0.230 | 0.230 | 0.024 | 0.011 | 5.88 | 2.94 | 1.18 |
| AARS | Glycyl-tRNA synthetase | GlyRS | 0.085 | 0.085 | 0.009 | 0.004 | 2.11 | 1.06 | 0.42 |
| AARS | Histidyl-tRNA synthetase | HisRS | 0.017 | 0.017 | 0.008 | 0.004 | 2.06 | 1.03 | 0.41 |
| AARS | Isoleucyl-tRNA synthetase | IleRS | 0.381 | 0.010 | 0.010 | 0.004 | 2.36 | 1.18 | 0.47 |
| AARS | Leucyl-tRNA synthetase | LeuRS | 0.041 | 0.008 | 0.008 | 0.004 | 2.02 | 1.01 | 0.40 |
| AARS | Lysyl-tRNA synthetase | LysRS | 0.109 | 0.109 | 0.027 | 0.012 | 6.74 | 3.37 | 1.35 |
| AARS | Methionine--tRNA synthetase | MetRS | 0.030 | 0.005 | 0.005 | 0.002 | 1.28 | 0.64 | 0.26 |
| AARS | Phenylalanyl-tRNA synthetase | PheRS | 0.135 | 0.135 | 0.007 | 0.003 | 1.67 | 0.84 | 0.33 |
| AARS | Prolyl-tRNA synthetase | ProRS | 0.154 | 0.006 | 0.006 | 0.003 | 1.53 | 0.77 | 0.31 |
| AARS | Seryl-tRNA synthetase | SerRS | 0.038 | 0.004 | 0.004 | 0.002 | 1.00 | 0.50 | 0.20 |
| AARS | Threonyl-tRNA synthetase | ThrRS | 0.082 | 0.082 | 0.008 | 0.004 | 1.96 | 0.98 | 0.39 |
| AARS | Tryptophanyl-tRNA synthetase | TrpRS | 0.163 | 0.163 | 0.016 | 0.007 | 3.86 | 1.93 | 0.77 |
| AARS | Tyrosyl-tRNA synthetase | TyrRS | 0.012 | 0.002 | 0.002 | 0.001 | 0.51 | 0.26 | 0.10 |
| AARS | Valyl-tRNA synthetase | ValRS | 0.016 | 0.016 | 0.005 | 0.002 | 1.13 | 0.57 | 0.23 |
| Translation Factors | Initiation factor 1 | IF1 | 0.104 | 0.104 | 0.026 | 0.012 | 6.43 | 3.21 | 1.29 |
| Translation Factors | Initiation factor 2 | IF2 | 0.041 | 0.041 | 0.010 | 0.005 | 2.51 | 1.26 | 0.50 |
| Translation Factors | Initiation factor 3 | IF3 | 0.456 | 0.456 | 0.228 | 0.102 | 56.52 | 28.26 | 11.30 |
| Translation Factors | Elongation factor G | EF-G | 0.636 | 0.636 | 0.318 | 0.142 | 78.85 | 39.43 | 15.77 |
| Translation Factors | Elongation factor Thermal unstable | EF-Tu | 11.274 | 11.274 | 2.255 | 1.007 | 559.34 | 279.67 | 111.87 |
| Translation Factors | Elongation factor Thermal stable | EF-Ts | 1.588 | 1.588 | 1.270 | 0.567 | 315.11 | 157.56 | 63.02 |
| Translation Factors | Release factor 1 | RF1 | 0.247 | 0.247 | 0.025 | 0.011 | 6.12 | 3.06 | 1.22 |
| Translation Factors | Release factor 2 | RF2 | 0.230 | 0.230 | 0.023 | 0.010 | 5.71 | 2.86 | 1.14 |
| Translation Factors | Release factor 3 | RF3 | 0.162 | 0.162 | 0.032 | 0.015 | 8.06 | 4.03 | 1.61 |
| Translation Factors | Ribosome recycling factor | RRF | 0.458 | 0.458 | 0.046 | 0.020 | 11.35 | 5.68 | 2.27 |
| Energy Regeneration | Methionyl-tRNA formyltransferase | MTF | 0.568 | 0.568 | 0.057 | 0.025 | 14.08 | 7.04 | 2.82 |
| Energy Regeneration | Creatine kinase | CK | 0.089 | 0.089 | 0.022 | 0.010 | 5.55 | 2.77 | 1.11 |
| Energy Regeneration | Adenylate kinase (Myokinase) | MK | 0.132 | 0.132 | 0.044 | 0.020 | 10.90 | 5.45 | 2.18 |
| Energy Regeneration | Nucleotide diphosphate kinase | NDK | 0.064 | 0.064 | 0.029 | 0.013 | 7.16 | 3.58 | 1.43 |
| Energy Regeneration | Inorganic pyrophosphatase | PPiase | 0.030 | 0.030 | 0.015 | 0.007 | 3.72 | 1.86 | 0.74 |
| T7 RNAP | T7 RNA polymerase | T7 RNAP | 0.100 | 0.100 | 0.050 | 0.022 | 12.37 | 6.19 | 2.47 |
| | | **Total** | **19.02** | **17.95** | **4.67** | **2.08** | **1157.53** | **578.77** | **231.51** |

Supplementary Table 3: PURE proteins concentration with T7 RNAP at full concentration across different versions.

| Category | PURE protein | | PURE v1 concentration (µg/mL) | PURE v2 concentration (µg/mL) | PURE v3 concentration (µg/mL) | PURE v3.90 concentration (µg/mL) | PURE v3.50 concentration (µg/mL) | PURE v3.25 concentration (µg/mL) | PURE v3.10 concentration (µg/mL) |
|---|---|---|---|---|---|---|---|---|---|
| AARS | Alanyl-tRNA synthetase | AlaRS | 70 | 70 | 2.00 | 0.89 | 0.50 | 0.25 | 0.10 |
| AARS | Arginyl-tRNA synthetase | ArgRS | 2 | 2 | 1.00 | 0.45 | 0.25 | 0.12 | 0.05 |
| AARS | Asparaginyl-tRNA synthetase | AsnRS | 22 | 1.1 | 1.10 | 0.49 | 0.27 | 0.14 | 0.05 |
| AARS | Aspartate-tRNA synthetase | AspRS | 8 | 8 | 0.40 | 0.18 | 0.10 | 0.05 | 0.02 |
| AARS | Cysteinyl-tRNA synthetase | CysRS | 1.2 | 0.1 | 0.10 | 0.04 | 0.02 | 0.01 | 0.00 |
| AARS | Glutaminyl-tRNA synthetase | GlnRS | 3.8 | 1 | 1.00 | 0.45 | 0.25 | 0.12 | 0.05 |
| AARS | Glutamyl-tRNA synthetase | GluRS | 12.6 | 12.6 | 1.30 | 0.58 | 0.32 | 0.16 | 0.06 |
| AARS | Glycyl-tRNA synthetase | GlyRS | 9.6 | 9.6 | 0.96 | 0.43 | 0.24 | 0.12 | 0.05 |
| AARS | Histidyl-tRNA synthetase | HisRS | 0.8 | 0.8 | 0.40 | 0.18 | 0.10 | 0.05 | 0.02 |
| AARS | Isoleucyl-tRNA synthetase | IleRS | 40 | 1 | 1.00 | 0.45 | 0.25 | 0.12 | 0.05 |
| AARS | Leucyl-tRNA synthetase | LeuRS | 4 | 0.8 | 0.80 | 0.36 | 0.20 | 0.10 | 0.04 |
| AARS | Lysyl-tRNA synthetase | LysRS | 6.4 | 6.4 | 1.60 | 0.71 | 0.40 | 0.20 | 0.08 |
| AARS | Methionine--tRNA synthetase | MetRS | 2.3 | 0.4 | 0.40 | 0.18 | 0.10 | 0.05 | 0.02 |
| AARS | Phenylalanyl-tRNA synthetase | PheRS | 17 | 17 | 0.85 | 0.38 | 0.21 | 0.11 | 0.04 |
| AARS | Prolyl-tRNA synthetase | ProRS | 10 | 0.4 | 0.40 | 0.18 | 0.10 | 0.05 | 0.02 |
| AARS | Seryl-tRNA synthetase | SerRS | 1.9 | 0.2 | 0.20 | 0.09 | 0.05 | 0.02 | 0.01 |
| AARS | Threonyl-tRNA synthetase | ThrRS | 6.2 | 6.2 | 0.60 | 0.27 | 0.15 | 0.07 | 0.03 |
| AARS | Tryptophanyl-tRNA synthetase | TrpRS | 6.3 | 6.3 | 0.60 | 0.27 | 0.15 | 0.07 | 0.03 |
| AARS | Tyrosyl-tRNA synthetase | TyrRS | 0.6 | 0.1 | 0.10 | 0.04 | 0.02 | 0.01 | 0.00 |
| AARS | Valyl-tRNA synthetase | ValRS | 1.8 | 1.8 | 0.50 | 0.22 | 0.12 | 0.06 | 0.02 |
| Translation Factors | Initiation factor 1 | IF1 | 1 | 1 | 0.25 | 0.11 | 0.06 | 0.03 | 0.01 |
| Translation Factors | Initiation factor 2 | IF2 | 4 | 4 | 1.00 | 0.45 | 0.25 | 0.12 | 0.05 |
| Translation Factors | Initiation factor 3 | IF3 | 10 | 10 | 5.00 | 2.23 | 1.24 | 0.62 | 0.25 |
| Translation Factors | Elongation factor G | EF-G | 50 | 50 | 25.00 | 11.16 | 6.20 | 3.10 | 1.24 |
| Translation Factors | Elongation factor Thermal unstable | EF-Tu | 500 | 500 | 100.00 | 44.65 | 24.81 | 12.40 | 4.96 |
| Translation Factors | Elongation factor Thermal stable | EF-Ts | 50 | 50 | 40.00 | 17.86 | 9.92 | 4.96 | 1.98 |
| Translation Factors | Release factor 1 | RF1 | 10 | 10 | 1.00 | 0.45 | 0.25 | 0.12 | 0.05 |
| Translation Factors | Release factor 2 | RF2 | 10 | 10 | 1.00 | 0.45 | 0.25 | 0.12 | 0.05 |
| Translation Factors | Release factor 3 | RF3 | 10 | 10 | 2.00 | 0.89 | 0.50 | 0.25 | 0.10 |
| Translation Factors | Ribosome recycling factor | RRF | 10 | 10 | 1.00 | 0.45 | 0.25 | 0.12 | 0.05 |
| Energy Regeneration | Methionyl-tRNA formyltransferase | MTF | 20 | 20 | 2.00 | 0.89 | 0.50 | 0.25 | 0.10 |
| Energy Regeneration | Creatine kinase | CK | 4 | 4 | 1.00 | 0.45 | 0.25 | 0.12 | 0.05 |
| Energy Regeneration | Adenylate kinase (Myokinase) | MK | 3 | 3 | 1.00 | 0.45 | 0.25 | 0.12 | 0.05 |
| Energy Regeneration | Nucleotide diphosphate kinase | NDK | 1.1 | 1.1 | 0.50 | 0.22 | 0.12 | 0.06 | 0.02 |
| Energy Regeneration | Inorganic pyrophosphatase | PPiase | 1 | 1 | 0.50 | 0.22 | 0.12 | 0.06 | 0.02 |
| T7 RNAP | T7 RNA polymerase | T7 RNAP | 10 | 10 | 10.00 | 10.00 | 10.00 | 10.00 | 10.00 |
| | | **Total** | **920.60** | **839.90** | **206.56** | **97.77** | **58.76** | **34.38** | **19.75** |

Supplementary Table 4: Buffers used for purification

**PURE protein purification buffers**

| Compound | Catalog number | Company | Buffer A | Buffer B | HT buffer | Stock buffer | Note |
|---|---|---|---|---|---|---|---|
| | | | mM | mM | mM | mM | |
| HEPES | H0887-100ML | Sigma-Aldrich | 50 | 50 | 50 | 50 | pH = 7.6 |
| Magnesium chloride | 63020-1L | Honeywell Fluka | 10 | 10 | 10 | 10 | |
| Ammonium chloride | 09718-250G | Sigma-Aldrich | 1000 | | | | |
| Potassium chloride | P5405-1KG | Sigma-Aldrich | | 100 | 100 | 100 | |
| Imidazole | I2399 | Sigma-Aldrich | | 500 | | | pH = 7.0 |
| Glycerol | G7757-1L | Sigma-Aldrich | | | | 30% | |
| β-mercaptoethanol | M6250-100ML | Sigma-Aldrich | 7 | 7 | 7 | 7 | |

**Ribosome purification buffers**

| Compound | Catalog number | Company | Lysis buffer | Elution buffer | Ribosome buffer | Note |
|---|---|---|---|---|---|---|
| | | | mM | mM | mM | |
| HEPES | H0887-100ML | Sigma-Aldrich | | | 20 | |
| Tris-HCl | BP152-500 | Fisher | 20 | 20 | | pH = 7.6 |
| Magnesium chloride | 63020-1L | Honeywell Fluka | 10 | 10 | | |
| Ammonium chloride | 09718-250G | Sigma-Aldrich | 30 | 30 | | |
| Imidazole | I2399 | Sigma-Aldrich | | 150 | | pH = 7.0 |
| Magnesium acetate | M0631 | Sigma-Aldrich | | | 6 | |
| Potassium chloride | P5405-1KG | Sigma-Aldrich | 150 | 150 | 30 | |
| β-mercaptoethanol | M6250-100ML | Sigma-Aldrich | 7 | 7 | 7 | |

Supplementary Table 5: Energy solution

| Component | Catalog number | Company | Concentration in energy solution (2.5x) | Final concentration in the reaction (1x) | Units |
|---|---|---|---|---|---|
| HEPES | H0887-100ML | Sigma-Aldrich | 125 | 50 | mM |
| Potassium glutamate | 49601 | Sigma-Aldrich | 250 | 100 | mM |
| Magnesium acetate | M0631 | Sigma-Aldrich | 29.5 | 11.8 | mM |
| Creatine phosphate | 27920 | Sigma-Aldrich | 50 | 20 | mM |
| TCEP | 646547 | Sigma-Aldrich | 2.5 | 1 | mM |
| Folinic acid | PHR1541 | Sigma-Aldrich | 0.05 | 0.02 | mM |
| Spermidine | S2626 | Sigma-Aldrich | 5 | 2 | mM |
| Amino Acid solution | LAA21-1KT | Sigma-Aldrich | 0.75 | 0.3 | mM |
| ATP | R0481 | ThermoFisher | 5 | 2 | mM |
| GTP | R0481 | ThermoFisher | 5 | 2 | mM |
| CTP | R0481 | ThermoFisher | 2.5 | 1 | mM |
| UTP | R0481 | ThermoFisher | 2.5 | 1 | mM |
| tRNA | 10109541001 | Roche | 130 | 52 | $U_{A260}$/mL |

Supplementary Table 6: eGFP DNA sequence

| | |
|---|---|
| eGFP linear DNA fragment | 5’-gatcttaaggctagagtactaatacgactcactatagggagaccacaacggtttccctctagaaataattttgtttaacttaagaaggaggaaaaaaaaATGTCTAAAGGTGAAGAATTATTCACTGGTGTTGTCCCAATTTTGGTTGAATTAGATGGTGATGTTAATGGTCACAAATTTTCTGTCTCCGGTGAAGGTGAAGGTGATGCTACTTACGGTAAATTGACCTTAAAATTTATTTGTACTACTGGTAAATTGCCAGTTCCATGGCCAACCTTAGTCACTACTTTAACTTATGGTGTTCAATGTTTTTCTAGATACCCAGATCATATGAAACAACATGACTTTTTCAAGTCTGCCATGCCAGAAGGTTATGTTCAAGAAAGAACTATTTTTTTCAAAGATGACGGTAACTACAAGACCAGAGCTGAAGTCAAGTTTGAAGGTGATACCTTAGTTAATAGAATCGAATTAAAAGGTATTGATTTTAAAGAAGATGGTAACATTTTAGGTCACAAATTGGAATACAACTATAACTCTCACAATGTTTACATCATGGCTGACAAACAAAAGAATGGTATCAAAGTTAACTTCAAAATTAGACACAACATTGAAGATGGTTCTGTTCAATTAGCTGACCATTATCAACAAAATACTCCAATTGGTGATGGTCCAGTCTTGTTACCAGACAACCATTACTTATCCACTCAATCTGCCTTATCCAAAGATCCAAACGAAAAGAGAGACCACATGGTCTTGTTAGAATTTGTTACTGCTGCTGGTATTACCCATGGTATGGATGAATTGTACAAATAAtaacgactcaggctgctacgcctgtgtactggaaaacaaaaccaaaacccaaaaaacaaaaaaactgagcccattggtatcgtggaaggactctatcaaaaaaaaaaaaaaaaaaaaaaaaaaaaaactagcataaccccttggggcctctaaacgggtcttgaggggttttttg-3’ |
| Forward amplification primer | 5’- gatcttaaggctagagtactaatacgactcactatagggagacc-3’ |
| Reverse amplification primer | 5’-caaaaaacccctcaagacccgtttagag-3’ |
| T7 promotor sequence | 5’-taatacgactcactatagg-3’ |
| RBS | 5’-aaggag-3’ |
| eGFP coding sequence | 5’-ATGTCTAAAGGTGAAGAATTATTCACTGGTGTTGTCCCAATTTTGGTTGAATTAGATGGTGATGTTAATGGTCACAAATTTTCTGTCTCCGGTGAAGGTGAAGGTGATGCTACTTACGGTAAATTGACCTTAAAATTTATTTGTACTACTGGTAAATTGCCAGTTCCATGGCCAACCTTAGTCACTACTTTAACTTATGGTGTTCAATGTTTTTCTAGATACCCAGATCATATGAAACAACATGACTTTTTCAAGTCTGCCATGCCAGAAGGTTATGTTCAAGAAAGAACTATTTTTTTCAAAGATGACGGTAACTACAAGACCAGAGCTGAAGTCAAGTTTGAAGGTGATACCTTAGTTAATAGAATCGAATTAAAAGGTATTGATTTTAAAGAAGATGGTAACATTTTAGGTCACAAATTGGAATACAACTATAACTCTCACAATGTTTACATCATGGCTGACAAACAAAAGAATGGTATCAAAGTTAACTTCAAAATTAGACACAACATTGAAGATGGTTCTGTTCAATTAGCTGACCATTATCAACAAAATACTCCAATTGGTGATGGTCCAGTCTTGTTACCAGACAACCATTACTTATCCACTCAATCTGCCTTATCCAAAGATCCAAACGAAAAGAGAGACCACATGGTCTTGTTAGAATTTGTTACTGCTGCTGGTATTACCCATGGTATGGATGAATTGTACAAATAA-3’ |
| T7 terminator sequence | 5’-tagcataaccccttggggcctctaaacgggtcttgaggggttttttg-3’ |
